# *In situ* maturated early-stage human induced pluripotent stem cell-derived cardiomyocytes improve cardiac function by enhancing segmentary contraction in infarcted rats

**DOI:** 10.1101/2021.03.09.434658

**Authors:** Diogo Biagi, Evelyn Thais Fantozzi, Julliana C Campos-Oliveira, Marcus Vinicius Naghetini, Antonio F. Ribeiro, Sirlene Rodrigues, Isabella Ogusuku, Rubia Vanderlinde, Michelle Lopes Araújo Christie, Debora B. Mello, Antonio C. Campos de Carvalho, Marcos Valadares, Estela Cruvinel, Rafael Dariolli

## Abstract

The scant ability of cardiomyocytes to proliferate makes heart regeneration one of the biggest challenges of science. Current therapies do not contemplate heart re-muscularization. In this scenario, stem cell-based approaches have been proposed to overcome the lack of regeneration. We hypothesize PluriCell hiPSC-derived cardiomyocytes (hiPSC-CMs) could enhance rat’s cardiac function after myocardial infarction (MI). Animals were subjected to permanent occlusion of the Left-Ventricle (LV) anterior descending coronary artery (LAD). Seven days after MI, Early-stage hiPSC-CMs were injected intramyocardially. Rats were subjected to Echocardiography pre- and post-treatment. Thirty days after injections, treated rats displayed 6.2% human cardiac grafts, which were characterized molecularly. Left ventricle ejection fraction (LVEF) was improved by 7.8% in cell-injected rats, while placebo controls showed an 18.2% deterioration. Also, cell-treated rats displayed a 92% and 56% increase in radial and circumferential strains, respectively. Human cardiac grafts maturate in situ, preserving proliferation with 10% Ki67 and 3% PHH3 positive nuclei. Grafts were perfused by host vasculature with no evidence for immune rejection nor ectopic tissue formations. Our findings support *PluriCell hiPSC-CMs* as an alternative therapy to treat MI. The next steps of preclinical development include efficacy studies in large animals on the path to clinical-grade regenerative therapy targeting human patients.

## 1. Introduction

Cardiovascular diseases, in particular myocardial infarction (MI), are the leading cause of morbi-mortality worldwide [1,2]. The adult heart displays a limited regenerative capacity due to cardiomyocytes’ scant proliferation ability [3,4]. Therapies currently available partially preserve the heart’s structure and function without any sign of muscle regeneration. In this scenario, patients with decompensated outcomes ultimately will end up in severe heart failure condition requiring heart transplantation [5].

Alternatively, stem cell-based therapies have been proposed to overcome the lack of heart re-muscularization within the available therapeutics [5]. Remarkably, multipotent stem cells isolated from adult tissues such as bone marrow [6–10] and adipose tissue [11,12] have demonstrated unsatisfactory heart regeneration performance. Conversely, human pluripotent stem cells such as embryonic stem cells (hESCs) [13] and induced pluripotent stem cells (hiPSCs) [14,15] are promising sources for generating cardiomyocytes (hPSC-CMs) *in vitro* [16,17]. Contractile hPSC-CMs are immature compared with adult cardiomyocytes [18–20] a double-edged characteristic for cell therapy. These cells display proliferation capacity, which is gradually lost as maturation level increases over time *in vitro* and *in vivo* [18,21,22]. Nevertheless, hPSC-CMs reaching adult cardiomyocyte level have never been reported yet.

Independently of this limitation, several groups demonstrated the transplantation of hPSC-CMs enhances cardiac function through human cardiac tissue engraftment [21,23,32–34,24–31]. Despite promising results, the perfect interaction between host and graft (e.g., the mitigation of arrythmias), satisfactory graft maturation levels over time *in vivo*, and other fundamental questions such as cell dose, initial age of cells for injection, delivery method, and the degree of importance of each of these factors for an extensive and sustained structural and functional improvement remain elusive.

In the last decades our group has worked to developing stem cell-based therapies to treat myocardial infarction building tools [35–38] and testing trending therapies [9,11,12,39,40] always aiming patients [6–8]. Adult stem cell approaches to treat MI have yet to yield relevant results clinically [41] and it is generally accepted that other approaches are needed [42]. Here, we demonstrated that PluriCell hiPSC-CMs enhanced the overall cardiac function on infarcted rats under immunosuppression. Treated rats had their cardiac function improved due to human cardiac cells’ efficient engraftment into the fibrotic tissue. Moreover, human cardiac grafts showed enhanced maturation *in situ*, preserving some degree of proliferation capacity. Furthermore, our grafts were vascularized by host vessels and did not elicit significant immune rejection. These findings support further studies in larger animal models to conquer the limitations inherently present in a rodent cardiovascular system with the aim to develop scalable and efficient regenerative stem cell-based therapies for humans.

## 2. Results

### 2.1. Descriptive analysis of mortality and echocardiography-based randomization

Forty-two female rats were used in this study. Thirty-eight animals were subjected to a surgical thoracotomy for MI-induction (33 MIs and 5 sham-operated rats), four rats were used as healthy controls. Notably, only one rat died during MI induction and two others within the first six days after the procedure (Table 1). From forty-eight hours after MI until the end of the protocol (37 days after MI), SHAM and MI-induced rats received two daily doses of cyclosporine A (CsA). CsA toxicity was assessed by monitoring animals’ body mass weekly. Body mass changes were similar between groups over time (Figure S1A). Six days after MI induction, 39 rats (4 CTRLs, 5 SHAMs, and 30 MIs) were subjected to baseline echocardiography to assess left ventricle (LV) cardiac function. Two infarcted animals died due to anesthetic overload during the echocardiography (Table 1). The LV ejection fraction (LVEF) of each animal was estimated. LVEF values were used to randomize infarcted animals into two balanced groups: **(1)** PSC (placebo received injections of Pro-survival cocktail supplemented vehicle, 14 animals) and **(2)** CELL (PSC vehicle + approx. 10 million hiPSC-CMs, 14 animals) (Table S1).

**Table 1:** Overall mortality and excluded animals by phase and group.

| Status | Phase | N# | % (from total) | Cause | Group | N# | % (from total) |
| --- | --- | --- | --- | --- | --- | --- | --- |
| **dead** | MI induction | 1 | 2.4% | irreversible fibrillation | MI | 1 | 2.4% |
|  | 1-6 days after MI induction (pre-echocardiography) | 2 | 4.8% | anesthetic overload | MI | 1 | 2.4% |
|  |  |  |  | sudden death * | MI | 1 | 2.4% |
|  | Baseline echocardiogram (day 6) | 2 | 4.8% | anesthetic overload | MI | 2 | 4.8% |
|  | Injection's procedure | 6 | 14.3% | cardiorespiratory arrest | PSC | 3 | 7.1% |
|  |  |  |  |  | CELL | 3 | 7.1% |
|  | 1-30 days after injection (pre-final echocardiography) | 6 | 14.3% | sudden death * | SHAM | 1 | 2.4% |
|  |  |  |  |  | PSC | 3 | 7.1% |
|  |  |  |  |  | CELL | 2 | 4.8% |
|  | Final echocardiogram (day 37) | 0 | 0.0% | - |  | 0 | 0.0% |
| **live** | LVEF cutoff (based on CTRLs) | 4 ** | 9.5% | 20% impairment versus CTRL animals at baseline | PSC | 2 | 4.8% |
|  |  |  |  |  | CELL | 2 | 4.8% |
|  | Completed follow-up & used for further analysis | 21 | 50.0% | - | CTRL | 4 | 9.5% |
|  |  |  |  | - | SHAM | 4 | 9.5% |
|  |  |  |  | - | PSC | 6 | 14.2% |
|  |  |  |  | - | CELL | 7 | 16.7% |
| **total** |  | 42 | 100% |  |  | 42 | 100% |

\* These animals were found dead in their cages. \*\* These animals were excluded from subsequent analysis.

Seven days after MI induction, PSC and CELL rats were subjected to a second surgical thoracotomy for cell transplantation (or placebo). Six rats (3 PSC and 3 CELL) died during the intramyocardial injection surgery (Table 1). The thirty-one remaining rats were followed 30 days after injections (Figure 1A). Throughout the four weeks, 1 SHAM animal died on day 19, 3 PSC rats died on days 14, 20, and 22, and 2 CELL animals died on days 16 and 26 after injection procedure (Table 1). Mortality rates were not significantly different between groups after treatment (Figure S1B). Finally, 2 PSC and 2 CELL rats were excluded from further comparative analyzes since they displayed less than 20% impairment of LVEF compared to healthy animals at baseline. Thus, all analyzes were performed comparing 4 CTRL, 4 SHAM, 6 PSC, and 7 CELL animals. The therapeutic protocol adopted was graphically represented in the Figure 1A.

**Figure 1.**
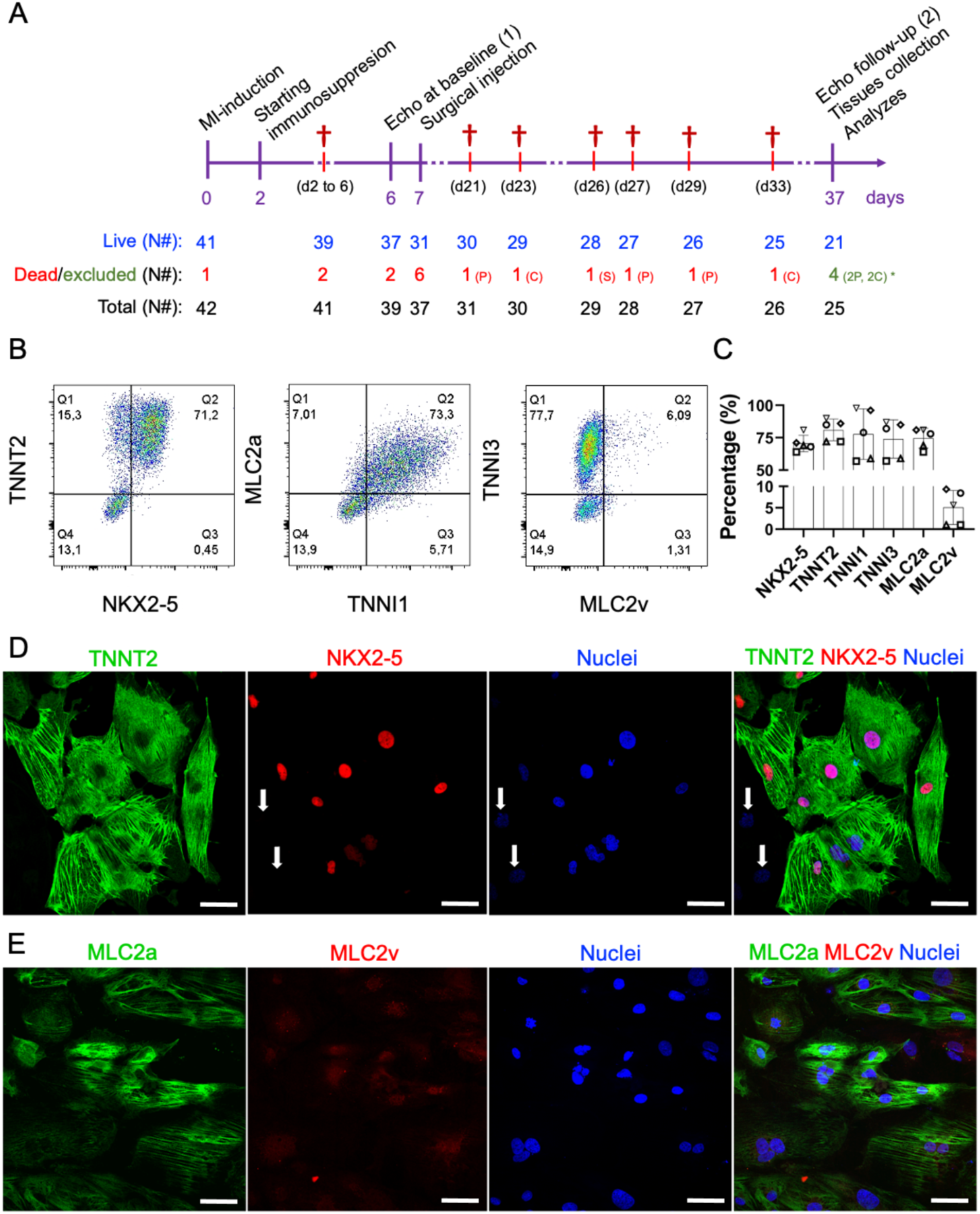
Therapeutic study design and molecular characterization of the early-stage PluriCell hiPSC-CMs. **A)** On day zero, myocardial infarctions were induced by permanent occlusion of the proximal-AD coronary artery. Forty-eight hours after MI (day 2), the immunosuppression protocol was started. CsA was administered twice a day intraperitoneally until the end of the protocol (on day 37). Six days after MI induction (day 6), all animals were subjected to baseline echocardiography and randomized. On day 7, animals from placebo (pro-survival cocktail - PSC) and CELL groups were subjected to a second surgical thoracotomy followed by intramyocardial injection of hiPSC-CMs. Thirty days after injections (on day 37), all survivors were subjected to final echocardiography and then were euthanized, necropsied, and had their hearts and other organs collected and stored for further analysis. **B)** Representative scatterplots of the injected hiPSC-CM population on day 11-15. **C)** The number of positive cells expressing cardiac markers such as TNNT2, TNNI1, TNNI3, MLC2A, MLC2V, and NKX2-5 was obtained by flow cytometry, expressed in percentage, and plotted in bar charts (mean ± SD). **D)** Representative immunofluorescence demonstrating the expression of cardiac markers in hiPSC-CMs on day 13 (TNNT2, NKX2-5 and Nuclei). Except for two cells in the image (white arrows), all the TNNT2 positive cells were also NKX2-5 positive. **E)** MLC2a, MLC2v, Nuclei, and Merged image. Note that the expression of MLC2v seems to be nuclear/perinuclear, strongly corroborating the lack of maturation of early-stage hiPSC-CMs. Scale bars: 100µm

### 2.2. Early-stage human iPSC-CMs are predominantly MLC2a positive (atrial-like cardiomyocytes) at the time of injection

Seven days after MI induction, 10 million early-stage hiPSC-CMs on the range of 11-15 days of differentiation were subjected to heat-shock priming [23,24], and intramyocardially injected (from epi-to myocardium) in 3 different points into the cardiac scar on MI-induced rats. Aliquots of the injections were collected and characterized by flow cytometry and immunofluorescence (Figure 1B-K). All animals received at least 70% of NK2 Homeobox 5 (NKX2-5) / Troponin T2 (TNNT2) positive cells (70.6% ± 5.7% and 81% ± 8.4%, Figure 1B-G). Furthermore, these cells expressed Troponin I1 (TNNI1, 77.8% ± 19.6) and Myosin light chain 2, atrial isoform (MLC2a, 74.6% ± 7.7%), both markers characteristic of early-stage hiPSC-CMs (Figure 1B-C). Despite their intrinsic immaturity, these cardiomyocytes also expressed Troponin I3 (TNNI3, 74% ± 14.7%) (Figure 1B-C). As expected, MLC2v (ventricular myosin light chain 2) was detected in a relatively small percentage of the population (5.1% ± 4%) (Figure 1B-C), also evidenced by immunofluorescence imaging (Figure 1H-K). Altogether, these findings demonstrated that all the animals were treated with a homogeneous population of hiPSC-CMs with low ranges of non-myocyte cells.

### 2.3. Early-stage human iPSC-CM therapy significantly improves the overall cardiac function of immunosuppressed infarcted rats

Four weeks after injections, grafts of human cardiac tissue were found in the LV of animals treated with hiPSC-CM therapy (Figure 2A). The mean percentage of the human graft was 6.23% ± 2.29% (human Ku80 (hKu80) / TNNT2 positive cells vs. fibrotic tissue, Figure 2B and Table S2). Interestingly, hiPSC-CM treated rats displayed significantly thicker MI-affected walls (* P<0.05 vs. PSC, Figure 2C, left bars). Conversely, we did not find any difference in thickness at free walls (Figure 2C, right bars). However, the percentage of injured area did not change on CELL group (vs. PSC rats, p=0.47, Figure S2A).

**Figure 2.**
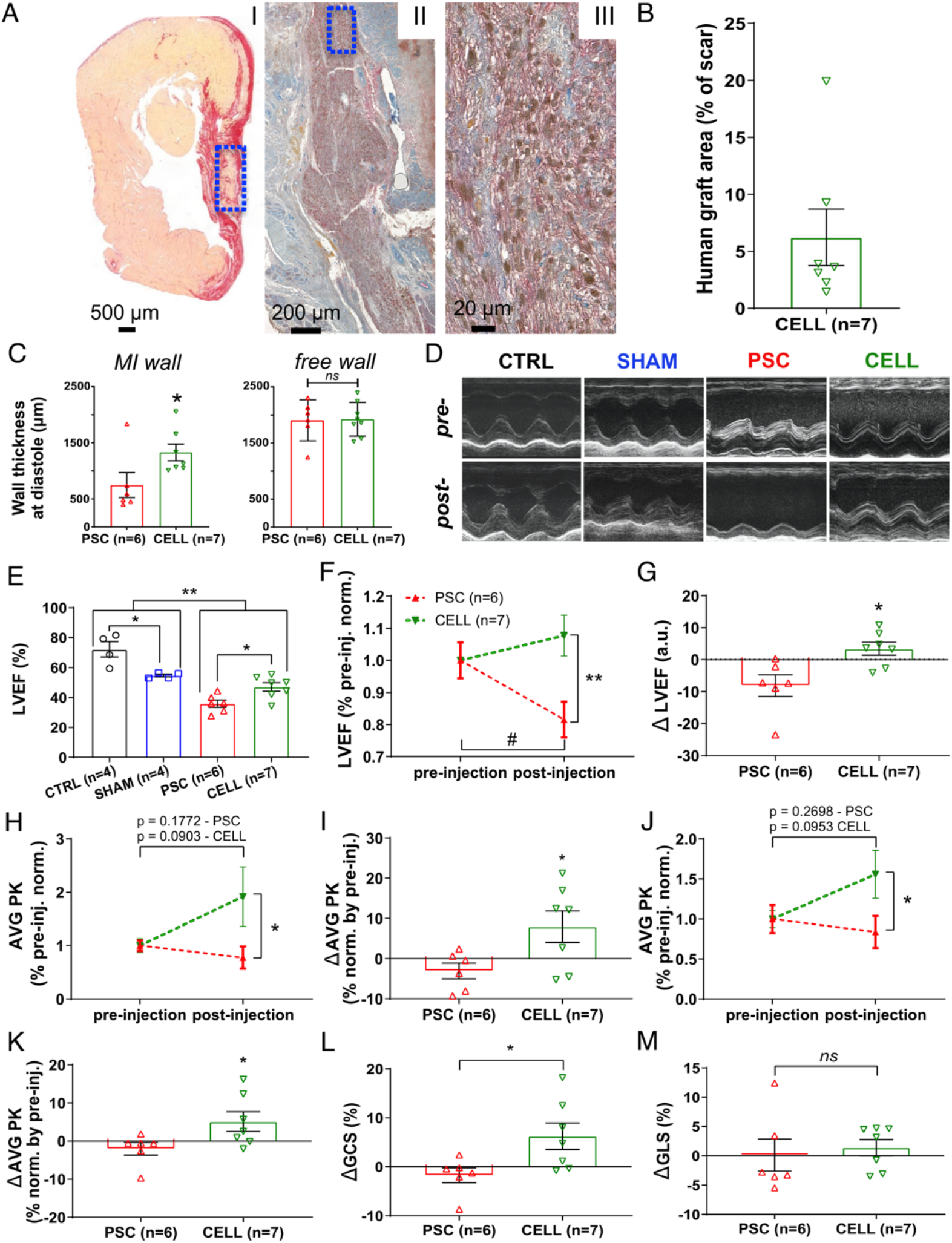
Early-stage hiPSC-CM-based therapy improves segmentary contraction resulting in overall LV cardiac function preservation on infarcted rats. Echocardiography was performed on days 6 (1-day prior injections) and 37 (30 days after injection) to evaluate the cell therapy effects. **A)** Representative panel of histological images from animal 29 (CELL group). **AI)** Papillary-level cross-section stained with picrosirius red evidencing “live” cardiac tissue (yellow) and fibrosis (red). “Live” cardiac tissue was found in the scar (Blue box). **AII)** Subsequent cross-section subjected to an immunohistochemical reaction against human Ku80 (brown nuclei) and Troponin T2 (red) evidencing human cells into the rat heart (Blue box). **AIII)** The human cardiac graft is taken from positive cells for both markers (brown nuclei – hKu80 and red fibers – Troponin T2). **B)** Percentage of the area covered by hKu80 positive cells into the scar tissue was quantified in 3-5 cross-sections at papillary level. The human graft areas were expressed in percentage relative to the scar area. All the CELL-treated rats displayed hKu80/Troponin T2 positive areas. **C)** Wall thickness at diastole was measured at papillary level using low magnification images. MI (left chart) and Free (right chart) wall values were calculated as a mean of three measurements covering the whole scar extension and expressed as mm. **D)** Representative short-axis of M-mode images pre-(day 6) and post-injection (day 37). **E)** LVEF was calculated using short-axis images at four LV deeps by a *Modified Simpson Algorithm*. CELL-treated rats showed improved LVEF 30 days post-treatment (vs. PSC, * P<0.05). Furthermore, MI-induced animals showed significantly lower LVEF than CTRL and SHAM groups (** p<0.01). Also, CTRL and SHAM displayed different LVEF (* p<0.05). **F)** LVEF of PSC and CELL rats were normalized by their respective pre-treatment values and plotted over time. CELL-treated rats show higher LVEF values than PSCs post-treatment (** p <0.01). PSCs showed a significant deterioration of LVEF 37 days after MI-induction (# p = 0.0145), whereas CELL rats had their LVEF preserved (p = 0.2162). **G)** LVEF delta reinforcing the cardiac function’s preservation on the CELL group (* p = 0.0117 vs. PSC). **H)** Radial Strain time-to-peak (AVG PK) of PSC and CELL rats were normalized by their respective pre-treatment values. CELL group display higher Radial strain AVG PK post-treatment (* p = 0.0441). Pre-vs. post-treatment analysis revealed no significant differences on PSC nor CELL group, as demonstrated by the p-values printed into the image. **I)** Radial Strain time-to-peak delta was also significantly different (* p <0.0371). **J)** Circumferential Strain time-to-peak (AVG PK) of PSC and CELL rats were normalized by their respective pre-treatment values. CELL group displays higher Circumferential strain AVG PK post-treatment (* p = 0.0494). Pre-vs. post-treatment analysis revealed no significant differences on PSC nor CELL group, as demonstrated by the p-values printed into the image. **K)** Circumferential Strain time-to-peak delta was also significantly different (* p = 0.0465). **L)** The Global Circumferential Strain delta was significantly positive for CELL-treated animals (p = 0.0324 vs. PSC), but **M)** The Global Longitudinal Strain delta did not display changes (data positively normalized; p = 0.6767 CELL vs. PSC). Data showed as mean ± SE.

Moreover, infarcted rats (PSC and CELL) showed significant impairment of their cardiac function at baseline, as evidenced by lower values of LVEF (vs. CTRL and SHAM animals, Figure S2B). In addition, PSC and CELL groups exhibited similar LVEF pre-treatment (Figure S2B). Nevertheless, 30 days after treatment, rats from the CELL group showed higher LVEF (vs. PSC group; Figure 2D-E). Indeed, most of the rats treated with placebo had significant deterioration of their cardiac function (Figure 2F; # p<0.05 vs. pre-treatment, and Figure S2C). Conversely, hiPSC-CM-treated animals had their LVEF preserved over time (Figure 2F; ** p<0.05 vs. PSC, and Figure S2D). The cell therapy’s benefit was evidenced through the positive LVEF delta calculated by pre-, and post-treatment LVEF values (* p<0.05, Figure 2G).

Besides the improvement in LVEF, the Radial (* p<0.05, Figure 2H) and Circumferential (* p<0.05, Figure 2J) strain time-to-peak (AVG PK), calculated by the average of 6 segments measured in short-axis images at papillary-level, were significantly higher on hiPSC-CM-treated rats than in the PSC group (and individual curves for each animal in Figure S3). In addition, both radial and circumferential strain time-to-peak deltas (* p<0.05, Figures 2I and K respectively) and global circumferential strain (Figure 2L) were significantly positive on CELL-treated animals.

Conversely, none of the previous parameters changed significantly for radial and longitudinal strain calculated using long-axis images (Figure 2M and Figure S4). Overall, the speckle tracking analysis corroborates LVEF improvement, strongly suggesting enhancements in segmentary contraction supporting active human graft presence into the mid-portion of the rat’s LV walls.

Despite the improvement in cardiac function observed in the CELL group, the LVEF of the treated rats is still significantly smaller than non-infarcted animals (CTRLs and SHAM) (Figure 2E).

### 2.4. The human graft is a cardiac tissue that preserves levels of cell cycling activity

Since the animals treated with early-stage hiPSC-CMs showed a significant improvement in their cardiac function, we assessed the human cardiac grafts cellular composition (Figure 3), cell cycle characteristics (Figure 4), and maturation levels (Figure 5 and 6).

**Figure 3.**
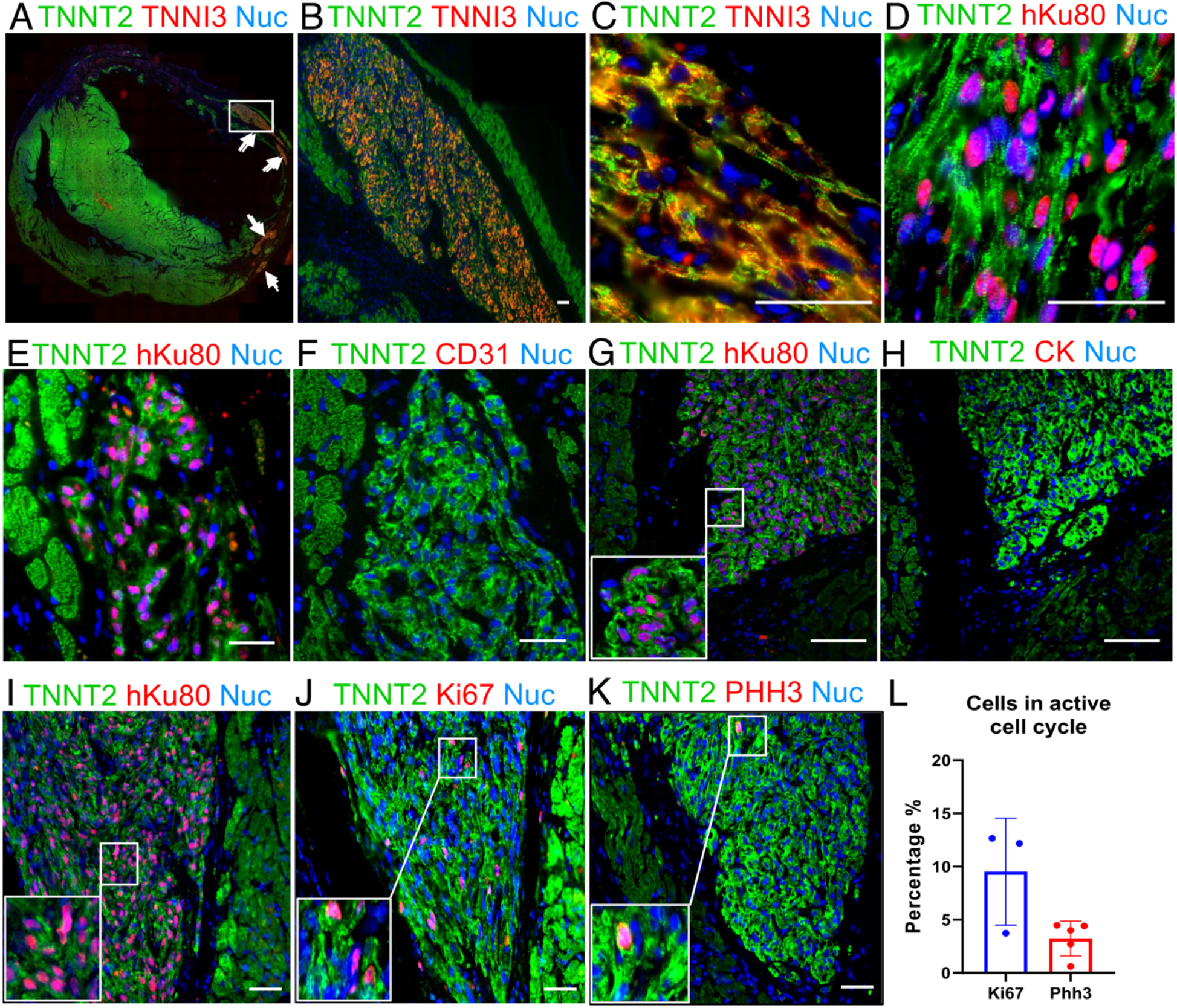
Representative image from human graft shown cardiac cellular composition. **A)** Panoramic immunofluorescence staining for TNNT2 (green) + human TNNI3 (Red), arrows are indicating grafts. **B)** Amplified view of largest graft region. **C)** Higher magnification of figure **B** showing sarcomere structure organization. **D)** Shows the evidence that hiPSC-CMs are following cardiac structure organization, TNNT2 (green) and hKu80 (pink) by overlap for hKu80 (red) and nuclei (blue). **E-H)** shows no evidence of hiPSC-CMs trans-differentiation into other cell types rather than cardiomyocytes. Subsequent slices were labeled for hKu80 **(E)** and human CD31 **(F)**. The same strategy was used to label hKu80 **(G)** and human Cytokeratin-1 **(H). I-L)** Ki67 and PHH3 markers evidenced that human cardiac grafts are composed by hiPSC-CMs with an active cell cycle. **I)** Subsequent slides were labeled by TNNT2 (green) and hKu80 (pink), **J)** TNNT2 (green) and Ki67 (pink), and **K)** TNNT2 (green) and PHH3 (pink). The color pink represents the overlap of red and blue (nuclei). **L)** Quantification for cell-cycle markers present a percentage of Ki67 positive cells and PHH3 positive cells (versus the total number of hKu80 positive nuclei for each image analyzed). Ki67: n=3 and PHH3: n=5. Scale bars: 50µm.

**Figure 4.**
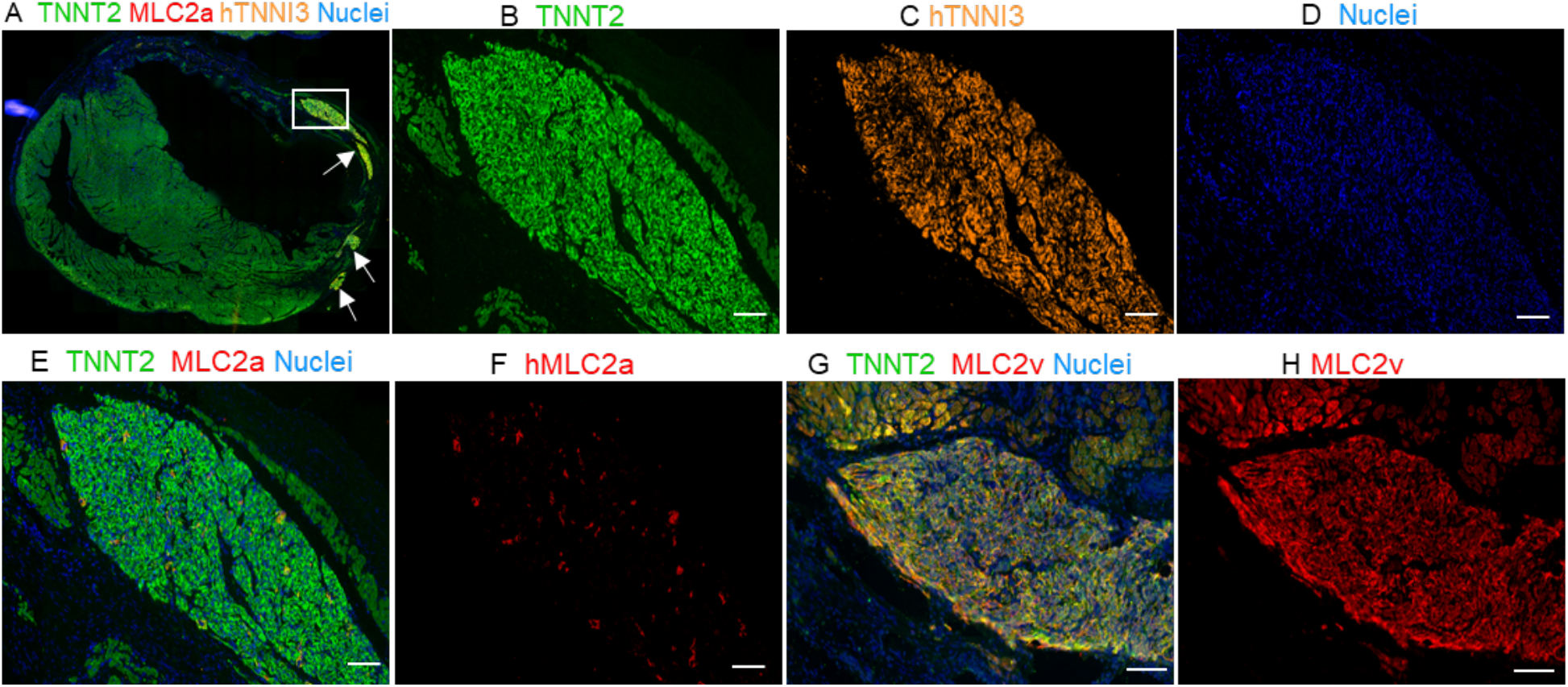
Immunofluorescence staining for hKu80 + MLC2v and MLC2a demonstrated maturation in the graft. **A)** Panoramic immunofluorescence staining for TNNT2 + human TNNI3. **B to D** shows a single channel for each marker. **B)** TNNT2 (green), **C)** human TNNI3 (orange) and **D)** nuclei (blue). **E)** Merged for TNNT2 (green), MLC2a (red) and nuclei (blue). **F)** Human MLC2a (red) showing poor labeling into the graft. **G)** Merged image for TNNT2 (green), MLC2v (red) and nuclei (blue). **H)** MLC2v showing massive labeling, compared with MLC2a. Scale bars: 100µm.

**Figure 5.**
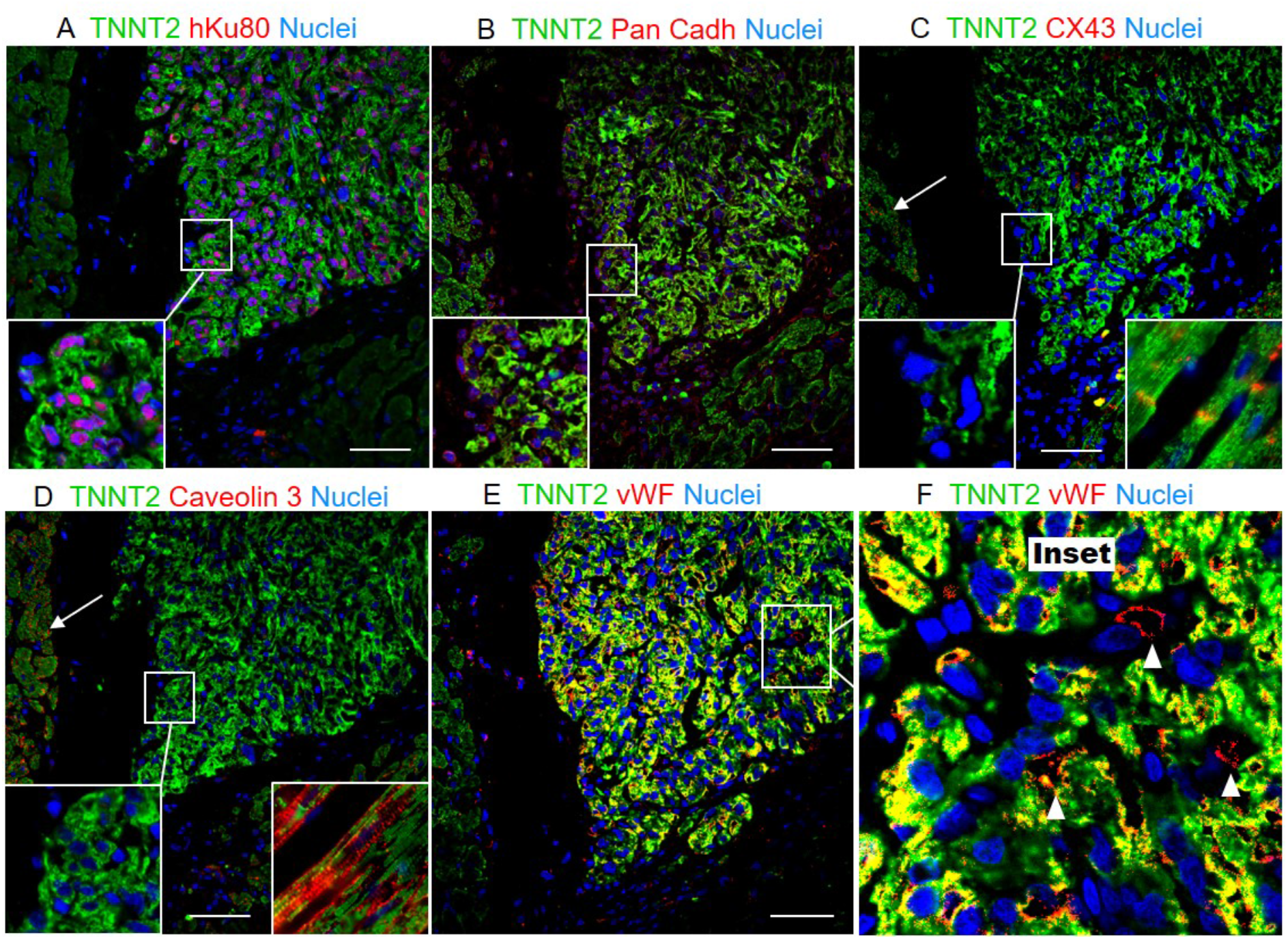
Histological evaluation of hiPSC-CMs myocardial grafts. **A)** Colocalization of hKu80 (red) and nuclei (blue), and TNNT2 (green). **B)** N-Cadherin expression represent a high cell-to-cell interaction. **C)** No evidence of Cx43 (red) nor **D)** Caveolin3 (red) was found in the grafted human tissues, suggesting hiPSC-CMs limited maturation. The amplified view evidenced insufficient labeling for Cx43 and Caveolin3 to be detectable into the grafts. Arrows shows Cx43 **(C)** and Caveolin 3 **(D)** labeled in the host cardiac tissue. The boxes show longitudinal sections in which the expected patterns of labeling for these proteins can be observed in the host tissue. **E-F)** Besides, rat vWF-positive blood vessels (red) perfusing the human cardiac grafts were observed in abundance in all the animals tested. **F)** Magnification of E, white arrow heads show individual vWF-positive vessels (red stain). Scale bars: 50µm.

**Figure 6.**
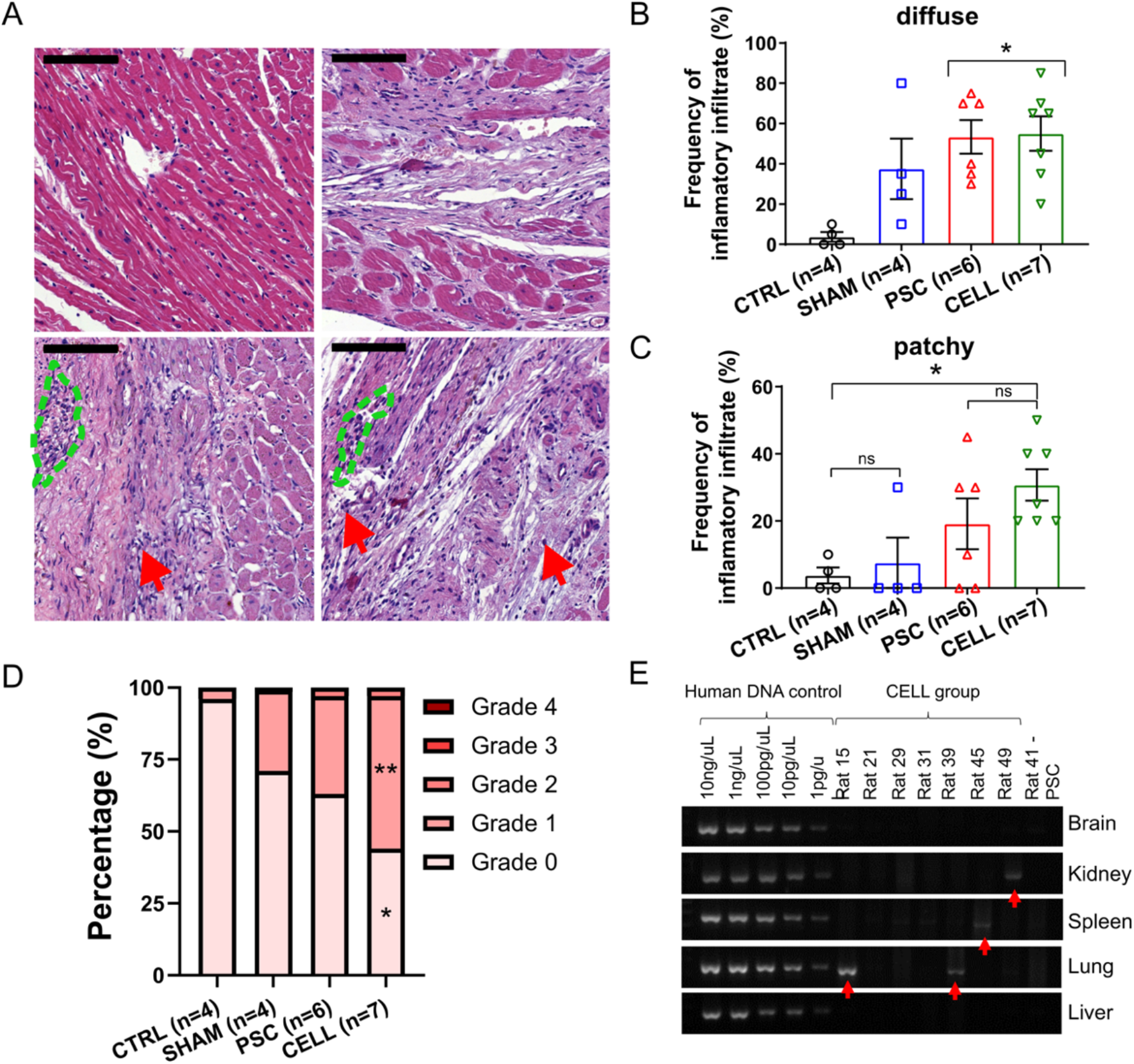
Immune-rejection against human cardiac grafts and the biodistribution of hiPSC-CMs over the body. **A)** pathological assessments were performed through H&E stained slides to qualify the incidence, distribution, and severity of inflammatory infiltrates. Representative images of heart sections for **A)** CTRL, SHAM, PSC and CELL rats (left to right, upper to bottom). Red arrows indicate sparse diffuse inflammatory cells, and the blue dashed forms indicate small patches of inflammatory cells (Scale bar: 100 µm). The frequency of **(B)** diffuse and **(C)** patchy infiltrates was qualitatively measured and expressed in the observation percentage. **D)** The severity of the inflammatory infiltrate was estimated using a well-established Grade system by observing the size and spread of inflammatory cells in the sections assessed. Grades were transformed in the percentage of severity per image and so per animal and plotted as stacked percentage bars. Grade 2 areas were observed only in infarcted animals. Grade 0 and 1 estimation were significantly different between CTRL and CELL groups (* p<0.05, and ** P<0.01, respectively). **E)** PCR results for human mitochondrial DNA indicate human material in rat 15 (lung), rat 39 (lung), rat 45 (spleen), and rat 49 (kidney) showed by the red arrows.

Overall, all the grafts identified were found in the myocardium, most of them in the scar area (Figure 3A). These human tissues displayed a highly compact structure and very distinct morphology compared to the host tissue (Figure 3B). Off notes, even using standard staining methods alone (e.g., Hematoxylin and Eosin - H&E), large grafts could be identified due to their distinct morphology. Besides the hKu80/TNNT2 expression (Figure 2A and Table 2), the human grafts massively expressed human TNNI3 (Figure 3A-C, Figure S5A and S5C). Also, structured sarcomeres were found in most of the grafts (Figure 3C-D and Figure S5B). Furthermore, we did not find evidence for human cells trans-differentiation into other cell types such as endothelial (human Platelet endothelial cell adhesion molecule (PECAM-1 or CD31) positive, Figure 3E-F and Figure S5D) or epithelial cells (human Cytokeratin-1 – CK, positive, Figure 3G-H and Figure S5E).

Adult cardiomyocytes display scant capacity of proliferation [43,44]. On the other hand, previous reports demonstrated that hiPSC-CMs could proliferate *in vitro* and *in vivo* [21,31]. Ki67 and Phosphohistone H3 (PHH3) are widespread proxies of active cell cycle [45]. We used sequential slides of hKu80 positive grafts (Figure 3I) to assess Ki67 (Figure 3J) and PHH3 (Figure 3K) expression into the human grafts. The percentage of Ki67 positive nuclei into the hKu80 grafts was 9.52% ± 5.02% (mean ± SD) (Figure 3J and L). In addition, the percentage of PHH3 positive nuclei into the hKu80 grafts was 3.24% ± 1.64% (mean ± SD) (Figure 3K and L).

### 2.5. The human cardiac grafts maturate in situ after injection

hiPSC-CMs are immature compared to adult cardiomyocytes [19,20]. Previous reports account for protein expression shifts (e.g., TNNI1 to TNNI3, MLC2a to MLC2v) to monitor growing levels of maturation of hiPSC-CMs *in vitro* and *in vivo* [22,31]. We used sequential slides of human TNNI3 (hTNNI3) / TNNT2 positive tissues (Figure 4A-D) to evaluate MLC2a (Figure 4E-F) and MLC2v (Figure 5G-H) expression in the human grafts. Approximately 30 days after the cell transplantation, the expression of MLC2a, highly expressed in the cells on days 11-15 (Figure 1H and K), was almost undetectable *in vivo* (Figure 4E-F and Figure S6A). Conversely, the expression of MLC2v was highly enhanced *in vivo* (Figure 4G-H vs. Figure 1I and K, and Figure S6B).

Additionally, the expression of N-Cadherin was identified in the human cardiac grafts, supporting cell-cell interactions in the grafted tissue (Figure 5A-B and Figure S6C). Despite the evidence of maturation and interactions between hiPSC-CMs *in vivo*, the expression of connexin 43 (Figure 5C) and Caveolin 3 (Figure 5D) proteins, other two crucial markers that support higher levels of cardiac-tissue maturation, were scant in the hKu80 positive grafts (Figure 5A, C-D and Figure S6D and S6E). Besides the expression of these markers, host-specific von Willebrand factor (vWF) positive blood vessels were found into the human cardiac grafts and surrounding boundaries, strongly suggesting an active perfusion of these grafts (Figure 5E-F and Figure S6F).

Altogether, the shift between MLC2a and MLC2v strongly suggests that **(1)** immature hiPSC-CMs injected matured over time *in vivo*; **(2)** the expression of N-Cadherin suggests active interaction between hiPSC-CMs composing the grafts, but **(3)** the scant expression of Connexin-43 (Cx43) and Caveolin-3 (Cav3) is strong evidence that the cells in the grafts, during the time frame of analysis, did not reach desired maturation levels to equalize the adult cardiac tissue. Furthermore, despite the limited maturation on the grafted tissues, **(4)** these xenografts were incorporated into the infarcted heart receiving perfusion from host blood vessels.

### 2.6. Immunosuppression confidently preserves human cardiac grafts from rejection or ectopic cellular formations

In addition to the presence of human cardiac tissue into infarcted hearts and the functional improvements observed in these rats, we evaluated whether inflammatory cells could surround human grafts as an additional metric to estimate human graft losses due to immunological rejection, despite active immunosuppressive protocol.

A histopathological analysis using H&E stained slides was performed to assess cellular inflammatory infiltrate incidence, distribution, and severity [12,46]. As expected, frequency of observation for inflammatory cells into the myocardium was high in both infarcted groups (PSC and CELL, Figure 6A-D). Notably, inflammatory infiltrates were also observed in SHAM animals, but in lower frequency than in infarcted rats (Figure 6A-D). Overall, the most frequent infiltrates were diffuse, where sparse cells can be observed solely or in very small groups (5-10 cells, Figure 6A red arrows), usually near to blood vessels (Figure 6A-B). Patches of cell infiltrates, where inflammatory cells can be seen in larger grouped, were also observed but in less frequency (Figure 6A, green-dashed areas, and C). Despite the incidence of inflammatory infiltrates, the degree of severity of these infiltrates was not significantly high (most of them classified as grades 0 and 1, Figure 6D). Statistical differences were observed between CTRL and CELL groups for the percentage of incidence of Grade 0 and 1 infiltrate, but no differences were observed comparing CELL-treated animals with SHAM or PSC rats (Figure 6D). Besides the histopathological assessment, we performed immunohistochemistry reactions using antibodies specific-against cluster of differentiation 3, 45 and 20 (CD3, CD45 and CD20, respectively) in 3-5 papillary-level cross-sections. As expected only scant diffuse cells or very small patches could be found into the heart of some of the SHAM (CD3: 2/4, CD45: 1/4, CD20: 2/3), PSC (CD3: 1/6, CD45: 2/6, CD20: 3/6) and CELL (CD3: 0/7, CD45: 5/7, CD20: 4/7) animals (Figure S7).

Finally, the biodistribution of hiPSC-CMs to the kidney, lung, spleen and brain, was evaluated to estimate harmful outcomes related to the presence of these cells out of their target organ and potential tumorigenic outcomes. Human mitochondrial DNA presence was assessed by a two-steps approach [34]. In the first screening using an end-point PCR, most of the samples were negative to human DNA contamination, except for lungs from animals 15 and 39, spleen from animal 45, and kidney from animal 49 (Figure 6E). The contaminated samples and two additional samples from different pieces of each tissue were subjected to a quantitative PCR. This analysis revealed minimal amounts of human DNA into two lung samples from animal 15 (0,02267% and 0,00195% human DNA to the total DNA, standard curve in the Figure S8). Additionally, no abnormal cell mass formations were found in these tissues (Figure S9).

These findings **(1)** corroborate the efficiency of the immunosuppression using cyclosporine A (CsA) and **(2)** suggest that the injection of hiPSC-CMs into the heart is safe since no abnormal formations nor **(3)** substantial human DNA were found in other organs rather than the heart.

## 3. Discussion

A rising number of pluripotent stem-cell-based therapies to treat myocardial infarction have been proposed. However, only marginal improvements in cardiac function have been reported from rodents to large animal models [47–49]. Here, we demonstrated **(1)** that the intramyocardial injection of 10 million early-stage hiPSC-CMs seven days after MI improved LV segmentary performance on infarcted rats. In addition, our findings show that **(2)** the human grafts massively expressed cardiomyocyte-related proteins, **(3)** with special emphases to the accentuated shift between MCL2a and MLC2v suggesting robust maturation *in situ*, **(4)** host/graft interactions and perfusion, **(5)** in the absence of substantial immune rejection.

Adult cardiomyocytes display a minimal regenerative capacity [43,44]. Conversely, hiPSC-CMs are immature compared to adult myocytes [19,20], preserving their capacity to proliferate *in vitro* and *in vivo* [21]. This ability is remarkably reduced over time as increasing levels of cellular maturation occur [21,31,50,51]. Furthermore, previous reports already demonstrated that larger human cardiac grafts were found into the heart of infarcted mice injected with day 20 cells than hiPSCs (day 0), mesoderm progenitors (day 4), cardiac progenitors (day 8), or later-stage hiPSC-CMs (day 30). Based on these previous findings and inspired by the successful work developed by Charles Murry’s group in the last decades [23–27,29,34], we intramyocardially injected PluriCell early-stage hiPSC-CMs (on days 11-15), heat-shock-primed [23], and diluted into a pro-survival cocktail [24] supplemented with GelTrex, seven days after the MI-induction.

In contrast to previous reports which injected cells in very acute (minutes-to-4 days after MI) [21,22,24,28,29,51] or chronic phases (4 -weeks later MI) [25] of the MI healing, here we tested hiPSC-CM therapy in a less inflammatory but still proliferative healing phase (7 days after MI). Endogenous repair mechanisms such as angiogenesis are active at this stage, favoring cells’ engraftment and viability [52–54]. On days 11-15, PluriCell hiPSC-CMs mainly expressed progenitor (e.g., NKX2-5) and early-stage cardiomyocyte markers (e.g., TNNI1, MLC2a) but did not express MLC2v, a proxy of more mature ventricular-like cells. Thirty days later, we found human cardiac grafts in all the injected rats averaging 6.23% ± 2.29% of the scar, with a maximum size of approximately 20%, twice more than demonstrated by *Laflamme et al. 2007* [24] and similar to *Fernandes et al. 2010* in a chronic model of ischemia/reperfusion [25].

Our rats were subjected to echocardiography pre- and 30 days post-injections. We demonstrated that human cardiac grafts positively affect the LV function of CELL-treated rats by increasing 7.8% LVEF at the end of the protocol. In contrast, the PSC group displayed 18.2% deterioration over time. Previous authors reported functional improvements using hESC-CMs and hiPSC-CMs acutely (minute-to-4 days after MI) in rodent models. *Laflamme et al*. demonstrated that the LV fractional shortening (LVSF) of rats treated with hESC-CMs was preserved four weeks after treatment [24]. *Fernandes et al*. showed an enhancement of approximately 28% on LVSF after a 4-week treatment of infarcted rats with hESC-CMs [29]. Furthermore, in mice models of MI, *Citro et al*. showed a 100% increase of LVSF and LVEF after four weeks of treatment compared to placebo [28], and *Saludas et al*. showed approximately 25% increase on LVEF 2 months after cell injection [51]. On the other hand, hESC-CM therapy to treat chronic MIs (injections 1-month after MI induction) seems ineffective in improving LV function in rodents [25].

Despite these exciting results, MI often affects specific heart segments resulting in hyperactivity of adjacent walls to compensate for muscle deterioration [43]. Measuring LV function simply by linear features may result in misinterpretation. All the reports discussed above provided echocardiographic results based on linear measurements. To avoid misinterpretation, we calculated LVEF using a *Modified Simpson Algorithm*, which considers four cross-sections of the LV to estimate volumetric changes pre-and post-treatment. Due to the higher quality of our analysis, we can conclude that our treatment resulted in a more realistic improvement of cardiac performance than previous reports.

In our study, human grafts were predominantly found in the lateral and anterior walls of LV, mostly at the papillary level. This observation prompted us to investigate whether the overall cardiac performance enhancement is associated with segmentary improvement due to human grafts. We assessed segmentary contraction by speckle tracking echocardiographic. Radial and circumferential strain calculated at the papillary-level in B-dimensional dynamic short-axis images were increased by 92% and 56%, respectively, in the CELL group. Conversely, PSC rats displayed 22% and 17% deterioration of those parameters. Nevertheless, radial, and longitudinal strain, which is calculated through long-axis images incorporating segments where human grafts were not found, did not change after treatment. Altogether, these data corroborate the segmentary improvement due to the presence of human cardiac grafts into the mid-portion of the myocardium on CELL-treated rats.

Additionally, we investigate the composition, proliferation capacity, maturation, and graft/host tissue interactions using immunofluorescence analysis. One month after the injection, PluriCell hiPSC-CM grafts identified by hKu80 or hTNNI3 expression solely displayed cardiac markers’ expression (e.g., TNNT2, TNNI3, MLC2v). Indeed, neither human CD31 nor cytokeratin positive cells were found in the grafts. Our data is corroborated by previous reports in different species [25,29,33,34]. Furthermore, we indirectly assess hiPSC-CM proliferation *in situ* by calculating the percentage of Ki67 and PHH3 positive cells in the grafts. These proxies are limited approximations which do not guarantee that cells in the active cell cycle will complete the cytokinesis [45]. Despite the technical limitation, approximately 10% of the hKu80 positive cells were also Ki67 positive (data obtained from 3 rats), and 3% were also PHH3 positive (data obtained from 5 rats), very similar to observed in previous reports [31,55]. In addition, *Funakoshi et al*. demonstrated *in vitro* that day 10 cells are twice more proliferative than day 20 and five times more proliferative than day 30 cells. These authors also tracked hiPSC-CM proliferation by Ki67 expression *in situ* up to 6 months after cell injection in a mice model of MI. hiPSC-CM proliferation was sharply reduced after three months of injection [21]. We believe that PluriCell hiPSC-CM-derived grafts might be growing *in vivo*. However, it is worth mentioning that, regardless of their proliferation capacity, the percentage of engrafted cells achieved in our study and others is still far from being translated into massive structural and functional improvements able to change myocardial infarction outcome completely.

Our grafts displayed remarkable maturation *in situ*, demonstrated by the shift between MCL2a and MLC2v expression. On days 11-15 (day of injections), our cells did not express MLC2v, a proxy for ventricular-like cell maturation. Thirty days later (41-45 days old cells), the expression of MLC2a was almost abolished, whereas MLC2v expression raised virtuously. Human grafts expressed substantial amounts of N-cadherin, but the expression of Cx43 and Cav3 was rarely observed compared to the host tissue. Furthermore, the human cardiac tissue was vascularized by the host blood vessels vWF positive. These data are corroborated by previous reports in different species [22,30,31,33,34]. Based on previous reports which assessed cardiac grafts up to 6 months after injection [21,22,30], we believe that more prolonged periods of post-treatment would also enhance the PluriCell hiPSC-CM grafts’ maturation *in vivo*.

Although we strove to work with clinically relevant conditions, it is worth listing few limitations to guide our result’s interpretation. Although PluriCell Biotech disposes of other well-characterized hiPSC lines in its portfolio, we used only one cell line showing superior reproducibility regarding cardiac differentiation [37]. We cannot exclude the possibility that other cell lines behave slightly differently in this context. On the other hand, future clinical applications might incorporate the concept of *off-the-shelf therapy* where a Universal non-immunogenic cell would be desirable. Second, a xenotransplant always imposes additional immunogenic complexities. We implemented a very standard cyclosporine A protocol to mitigate human cells’ rejection. Blood samples for dosing CsA were collected during the study. Due to transportation issues, these samples were lost, precluding an acceptable dosage of CsA over time. CsA can affect cardiac function through various mechanisms [56,57]. One of our SHAM animals suddenly died with clear signs of intoxication (e.g., cachexia, necrosis of abdominal skin and muscle). Also, the SHAM group displayed impaired LVEF compared to healthy rats (which did not receive immunosuppressants). We cannot exclude that survivors also had some degree of CsA-related compromising affecting overall cardiac function. In addition, our histopathological evaluation of immune rejection shows slight trends towards higher inflammatory infiltration into the heart of CELL-treated rats. Others similarly report this trend in different species and treatments [12,27,30,33,58,59].

Notably, our rats were exclusively treated with hiPSC-CMs without concurrent well-established drug-treatments such as beta-blockers and angiotensin-converting enzyme inhibitors, which can be a beneficial combination to use with a cell-therapy [12]. Importantly, our 30 days follow-up is short to observe more sustained improvements in cardiac function and graft maturation. Finally, based on the previous well-established reports regarding graft-related arrhythmias [26,27,30,31,33,34] and the rat model’s intrinsic limitations to access such a complex interplay between host and graft, we deliberately avoid assessing functional coupling of PluriCell hiPSC-CM grafts in this study. To overcome this limitation, we are already working with pig models of MI. We will study the cell’s coupling with the host tissue in this model that displays anatomopathological, metabolic, and even cellular conditions roughly similar to humans [12,32,33].

Altogether, our findings support the use of PluriCell early-stage hiPSC-CMs as an alternative therapy to regenerate segments of the myocardium on infarcted rats. The present work demonstrated that day 11-15 cells, injected seven days after the MI induction, **(1)** ameliorate overall cardiac function through segmental contraction enhancement; **(2)** display clear signs of maturation *in vivo* toward a ventricular-like phenotype **(3)** preserving some proliferation capacity. Moreover, our human grafts expressed interactive proteins corroborating the possibility of host/graft interaction. Finally, **(4)** we found the host circulation into human grafts in all the animals, and **(5)** we did not find significant immune rejection response nor abnormal tissue formations in or out of the heart. This proof-of-concept article is an additional resource in the endeavor towards developing a clinical-grade regenerative cell-based therapy to treat patients affected by degenerative cardiac diseases such as myocardial infarction.

## 4. Experimental Procedures

All procedures and analyses were carried by operators ***<u>blinded for experimental groups</u>***. Codes were revealed only when all the analyses were completed. Further detailed information regarding the following experimental procedures and materials is presented in a Supplementary Information document.

### 4.1. Ethics statement and animal care

This investigation agrees with the Declaration of Helsinki and the study protocol was approved by the Ethics Committee for Medical Research on Human Beings of the Institute of Biomedical Sciences from the University of Sao Paulo (#2.009.562) and the Committee on Animal Research and Ethics (CARE) at the Federal University of Rio de Janeiro (UFRJ) protocol #117/18. Signed informed consent was obtained from the cell donor. Animal experimentation and care agrees with the ARRIVE (Animals in Research: Reporting In Vivo Experiments) [60].

### 4.2. hiPSC-CM differentiation

Early-stage hiPSC-CMs (on days 11-15) were derived from the ACP5 hiPSC (a Pluricell Biotech hiPSC line). Briefly, hiPSCs were generated from healthy-donor erythroblasts using an episomal reprogramming system [37]. Cardiomyocytes were differentiated using a monolayer-based protocol previously described [37].

### 4.3. hiPSC-CM characterization: Flow cytometry and immunofluorescence

hiPSC-CMs were characterized by flow cytometry and immunofluorescence assays. Detailed lists of antibodies (Table S3), reagents (Table S4), and descriptive protocols are in the Supplementary information document.

### 4.4. Myocardial infarction induction

Two-month-old adult female Wistar rats (150-200g body weight) were subjected to surgical procedure to induce myocardial infarction (day 0). The left anterior descending coronary artery (LAD) was permanently occluded as previously described [38].

### 4.5. Immunosuppression

Except for the healthy CTRL animals, rats were treated with 20mg/kg of Cyclosporine A (Sandimmun IV, Novartis, Switzerland) [61–65] daily (2 doses of 10mg/kg every 12 hours), via intraperitoneal injection from day 2 to euthanasia.

### 4.6. Early-stage hiPSC-CM priming and intramyocardial injection

hiPSC-CMs were primed by Heat-Shock [23] before the intramyocardial injection. On day 7 of the experimental timeline, 10 million cells (on days 11-15) were suspended in 150uL of a pro-survival solution supplemented with approximately 0.4mg of GelTrex (ThermoFisher) [24] and directly injected into the infarcted rats’ left ventricle through a second surgical thoracotomy. Injections were split into three sites inside of and surrounding the scar tissue. Placebo animals were injected with pro-survival solution supplemented with GelTrex without cells.

### 4.7. Echocardiography, randomization, and exclusion criteria

Animals under inhalation anesthesia were subjected to echocardiography on days 6 (pre-) and 37 (post-injection). Parasternal images were captured using a Vevo® 2100 ultrasound equipment (VisualSonics, Canada) using a 25 MHz transducer. Analyses were performed using VevoCQ™ LV Analysis and Vevo LAB - Strain Analysis (VisualSonics, Canada). American Society of Echocardiography recommendations [66] were followed to data analysis and interpretation. On day 6, MI-induced rats were randomized by LVEF in two balanced groups. Moreover, animals (placebo or treated) with less than 20% LVEF impairment (at baseline) compared to control animals (LVEF above 55%) were excluded from the final analyses (functional and histopathological).

### 4.8. Euthanasia and tissues’ sampling

On day 37, after the final echocardiography after deep anesthesia, animals were euthanized by potassium chloride overdose. Thoracic and abdominal regions were photographed and the organs of interest (lung, liver, kidney, spleen and left cerebral hemisphere) were collected and stored at –80°C. Hearts were fixed using 4% paraformaldehyde (PFA) for further histopathological and molecular analyses.

### 4.9. Histology, immunohistochemistry (IHC) and immunofluorescent (IF) assays

The hearth was removed from the thoracic region after euthanasia. The tissue was cross-sectioned in the middle region, washed with 1x PBS, and PFA-fixed for histological, immunohistochemistry (IHC) and immunofluorescent (IF) staining (antibodies are listed in table S1). Details are in the Supplemental material. Immunofluorescence and immunohistochemistry micrographs were obtained using TissueFAXS slide scanner (TissueGnostics, Austria), confocal microscope LSM 800 (Zeiss-Germany) and a conventional fluorescence microscope Axio Imager 2 (Zeiss-Germany). The analyses were performed using ImageJ.

### 4.10. Biodistribution

To evaluate hiPSC-CMs migration and survival in other tissues after injection in the heart human mitochondrial DNA were searched in kidney, lung, brain, liver, and spleen using RT-PCR and RT-qPCR as previously described by *Liu et al*. [34].

### 4.11. Statistical analysis

Results were expressed as mean ± standard error of the mean, expect for different indication. One- or two-way analysis of variance (ANOVA) with Bonferroni *post-hoc* test or unpaired Student’s t-test were utilized to compare groups as appropriate. All statistical analyses were performed using GraphPad Prism 8.0 (GraphPad Software Inc., CA, USA). P-values < 0.05 were considered significant. For p<0.05 = *; p<0.001 = **; p<0.0005 = *** and p<0.0001 = ****, expect for different indication.

## Supplementary Materials

Supplementary methods, Tables and Figures were incorporated into a single DOCX file.

### Supplementary Tables

Table S1: LVEF distribution in MI-induced rats after randomization

Table S2: Percentage of hKu80 positive versus scar tissue area per animal

Table S3: Antibodies list

Table S4: List of reagents used in this study

### Supplementary Figures

Figure S1: Body weight control and survival rate

Figure S2: MI area and LVEF pre-injections

Figure S3: Short Axis Strain Analysis

Figure S4: Long Axis Strain Analysis

Figure S5: Morphology of the human graft

Figure S6: Maturation and cell-cell interaction

Figure S7: Immunohistochemistry for inflammatory markers

Figure S8: Human mitochondrial DNA were identified in some tissues

Figure S9: Necropsy

## Acknowledgments

We thank Dr. Talita Glaser and Prof. Alexander Henning Ulrich for assisting in using *TissueFAXS* equipment from Ulrich’s Laboratory at University of Sao Paulo.

## Author Contributions

Conceptualization, D.B, A.C.C.C, M.V, E.C and R.D; Methodology, D.B, E.T.F, J.C.C.O, M.V.N, A.F.R.Jr, M.L.A.C, D.B.M, A.C.C.C, M.V, E.C and R.D; Validation, D.B, M.V, E.C and R.D; Formal Analysis, D.B, E.C and R.D; Investigation, D.B, E.T.F, J.C.C.O, M.V.N, A.F.R.Jr, M.L.A.C, D.B.M, E.C and R.D; Resources, A.C.C.C, M.V and R.D; Data Curation, R.D; Writing – Original Draft Preparation, J.C.C.O, A.F.R.Jr., E.C and R.D; Writing – Review & Editing, D.B, E.T.F, J.C.C.O, M.V.N, A.F.R.Jr, S.R, I.O, R.V, M.L.A.C, D.B.M, A.C.C.C, M.V, E.C and R.D; Visualization, R.D; Supervision, R.D; Project Administration, R.D; Funding Acquisition, D.B, E.C, M.V and R.D. All authors have read and agreed to the published version of the manuscript.

## Institutional Review Board Statement

This investigation agrees with the Declaration of Helsinki and the study protocol was approved by the Ethics Committee for Medical Research on Human Beings of the Institute of Biomedical Sciences from the University of Sao Paulo (#2.009.562) and the Committee on Animal Research and Ethics (CARE) at the Federal University of Rio de Janeiro (UFRJ) protocol #117/18. Signed informed consent was obtained from the cell donor. Animal experimentation and care agrees with the ARRIVE (Animals in Research: Reporting In Vivo Experiments).

## Informed Consent Statement

The informed consent was obtained from the subject that donates blood samples for erythroblasts reprogramming into iPSCs in 2016.

## References

1. Teo, K.K.; Rafiq, T. Cardiovascular risk factors and prevention: a perspective from developing countries. Can. J. Cardiol. 2021, doi:10.1016/j.cjca.2021.02.009.

2. Virani, S.S.; Alonso, A.; Aparicio, H.J.; Benjamin, E.J.; Bittencourt, M.S.; Callaway, C.W.; Carson, A.P.; Chamberlain, A.M.; Cheng, S.; Delling, F.N.; et al. Heart Disease and Stroke Statistics—2021 Update. Circulation 2021, 143, doi:10.1161/CIR.0000000000000950.

3. Porrello, E.R.; Mahmoud, A.I.; Simpson, E.; Hill, J.A.; Richardson, J.A.; Olson, E.N.; Sadek, H.A. Transient regenerative potential of the neonatal mouse heart. Science 2011, 331, 1078–80, doi:10.1126/science.1200708.

4. Vivien, C.J.; Hudson, J.E.; Porrello, E.R. Evolution, comparative biology and ontogeny of vertebrate heart regeneration. npj Regen. Med. 2016, 1, 16012, doi:10.1038/npjregenmed.2016.12.

5. Mummery, C.L.; Davis, R.P.; Krieger, J.E. Challenges in using stem cells for cardiac repair. Sci. Transl. Med. 2010, 2, 27ps17, doi:10.1126/scitranslmed.3000558.

6. Gowdak, L.H.W.; Schettert, I.T.; Baptista, E.; Lopes, N.L.G.; Rochitte, C.E.; Vieira, M.L.C.; Grupi, C.J.; César, L.A.M.; Krieger, J.E.; de Oliveira, S. a Intramyocardial injection of autologous bone marrow cells as an adjunctive therapy to incomplete myocardial revascularization--safety issues. Clinics (Sao Paulo). 2008, 63, 207–14.

7. Gowdak, L.H.W.; Schettert, I.T.; Rochitte, C.E.; Lisboa, L.A.F.; Dallan, L.A.O.; César, L.A.M.; de Oliveira, S.A.; Krieger, J.E. Early Increase in Myocardial Perfusion After Stem Cell Therapy in Patients Undergoing Incomplete Coronary Artery Bypass Surgery. J. Cardiovasc. Transl. Res. 2011, 4, 106–113, doi:10.1007/s12265-010-9234-2.

8. Nicolau, J.C.; Furtado, R.H.M.; Silva, S.A.; Rochitte, C.E.; Rassi, A.; Moraes, J.B.M.C.; Quintella, E.; Costantini, C.R.; Korman, A.P.M.; Mattos, M.A.; et al. Stem-cell therapy in ST-segment elevation myocardial infarction with reduced ejection fraction: A multicenter, double-blind randomized trial. Clin. Cardiol. 2018, 41, 392–399, doi:10.1002/clc.22882.

9. Nakamuta, J.S.; Danoviz, M.E.; Marques, F.L.N.; dos Santos, L.; Becker, C.; Gonçalves, G. a; Vassallo, P.F.; Schettert, I.T.; Tucci, P.J.F.; Krieger, J.E. Cell therapy attenuates cardiac dysfunction post myocardial infarction: effect of timing, routes of injection and a fibrin scaffold. PLoS One 2009, 4, e6005, doi:10.1371/journal.pone.0006005.

10. dos Santos, L.; Santos, A. a; Gonçalves, G.; Krieger, J.E.; Tucci, P.J.F. Bone marrow cell therapy prevents infarct expansion and improves border zone remodeling after coronary occlusion in rats. Int. J. Cardiol. 2010, 145, 34–9, doi:10.1016/j.ijcard.2009.06.008.

11. Danoviz, M.E.; Nakamuta, J.S.; Marques, F.L.N.; dos Santos, L.; Alvarenga, E.C.; dos Santos, A. a; Antonio, E.L.; Schettert, I.T.; Tucci, P.J.; Krieger, J.E. Rat adipose tissue-derived stem cells transplantation attenuates cardiac dysfunction post infarction and biopolymers enhance cell retention. PLoS One 2010, 5, e12077, doi:10.1371/journal.pone.0012077.

12. Dariolli, R.; Naghetini, M. V; Marques, E.F.; Takimura, C.K.; Jensen, L.S.; Kiers, B.; Tsutsui, J.M.; Mathias, W.; Lemos Neto, P.A.; Krieger, J.E. Allogeneic pASC transplantation in humanized pigs attenuates cardiac remodeling post-myocardial infarction. PLoS One 2017, 12, e0176412, doi:10.1371/journal.pone.0176412.

13. Thomson, J.A. Embryonic Stem Cell Lines Derived from Human Blastocysts. Science (80-.). 1998, 282, 1145–1147, doi:10.1126/science.282.5391.1145.

14. Takahashi, K.; Tanabe, K.; Ohnuki, M.; Narita, M.; Ichisaka, T.; Tomoda, K.; Yamanaka, S. Induction of pluripotent stem cells from adult human fibroblasts by defined factors. Cell 2007, 131, 861–72, doi:10.1016/j.cell.2007.11.019.

15. Takahashi, K.; Yamanaka, S. Induction of pluripotent stem cells from mouse embryonic and adult fibroblast cultures by defined factors. Cell 2006, 126, 663–76, doi:10.1016/j.cell.2006.07.024.

16. Burridge, P.W.; Keller, G.; Gold, J.D.; Wu, J.C. Production of de novo cardiomyocytes: Human pluripotent stem cell differentiation and direct reprogramming. Cell Stem Cell 2012, 10, 16–28, doi:10.1016/j.stem.2011.12.013.

17. Lian, X.; Hsiao, C.; Wilson, G.; Zhu, K.; Hazeltine, L.B.; Azarin, S.M.; Raval, K.K.; Zhang, J.; Kamp, T.J.; Palecek, S.P. Robust cardiomyocyte differentiation from human pluripotent stem cells via temporal modulation of canonical Wnt signaling. Proc. Natl. Acad. Sci. U. S. A. 2012, 109, E1848–57, doi:10.1073/pnas.1200250109.

18. Babiarz, J.E.; Ravon, M.; Sridhar, S.; Ravindran, P.; Swanson, B.; Bitter, H.; Weiser, T.; Chiao, E.; Certa, U.; Kolaja, K.L. Determination of the human cardiomyocyte mRNA and miRNA differentiation network by fine-scale profiling. Stem Cells Dev. 2012, 21, 1956– 65, doi:10.1089/scd.2011.0357.

19. Lundy, S.D.; Zhu, W.-Z.; Regnier, M.; Laflamme, M.A. Structural and functional maturation of cardiomyocytes derived from human pluripotent stem cells. Stem Cells Dev. 2013, 22, 1991–2002, doi:10.1089/scd.2012.0490.

20. Robertson, C.; Tran, D.D.; George, S.C. Concise review: Maturation phases of human pluripotent stem cell-derived cardiomyocytes. Stem Cells 2013, 31, 829–837, doi:10.1002/stem.1331.

21. Funakoshi, S.; Miki, K.; Takaki, T.; Okubo, C.; Hatani, T.; Chonabayashi, K.; Nishikawa, M.; Takei, I.; Oishi, A.; Narita, M.; et al. Enhanced engraftment, proliferation, and therapeutic potential in heart using optimized human iPSC-derived cardiomyocytes. Sci. Rep. 2016, 6, 19111, doi:10.1038/srep19111.

22. Kadota, S.; Pabon, L.; Reinecke, H.; Murry, C.E. In Vivo Maturation of Human Induced Pluripotent Stem Cell-Derived Cardiomyocytes in Neonatal and Adult Rat Hearts. Stem Cell Reports 2017, 8, 278–289, doi:10.1016/j.stemcr.2016.10.009.

23. Laflamme, M.A.; Gold, J.; Xu, C.; Hassanipour, M.; Rosler, E.; Police, S.; Muskheli, V.; Murry, C.E. Formation of human myocardium in the rat heart from human embryonic stem cells. Am. J. Pathol. 2005, 167, 663–71, doi:10.1016/S0002-9440(10)62041-X.

24. Laflamme, M.A.; Chen, K.Y.; Naumova, A. V; Muskheli, V.; Fugate, J.A.; Dupras, S.K.; Reinecke, H.; Xu, C.; Hassanipour, M.; Police, S.; et al. Cardiomyocytes derived from human embryonic stem cells in pro-survival factors enhance function of infarcted rat hearts. Nat. Biotechnol. 2007, 25, 1015–1024, doi:10.1038/nbt1327.

25. Fernandes, S.; Naumova, a V; Zhu, W.Z.; Laflamme, M. a; Gold, J.; Murry, C.E. Human embryonic stem cell-derived cardiomyocytes engraft but do not alter cardiac remodeling after chronic infarction in rats. J. Mol. Cell. Cardiol. 2010, 49, 941–9, doi:10.1016/j.yjmcc.2010.09.008.

26. Shiba, Y.; Fernandes, S.; Zhu, W.Z.; Filice, D.; Muskheli, V.; Kim, J.; Palpant, N.J.; Gantz, J.; Moyes, K.W.; Reinecke, H.; et al. Human ES-cell-derived cardiomyocytes electrically couple and suppress arrhythmias in injured hearts. Nature 2012, 489, 322–325, doi:10.1038/nature11317.

27. Chong, J.J.H.; Yang, X.; Don, C.W.; Minami, E.; Liu, Y.-W.; Weyers, J.J.; Mahoney, W.M.; Van Biber, B.; Cook, S.M.; Palpant, N.J.; et al. Human embryonic-stem-cell-derived cardiomyocytes regenerate non-human primate hearts. Nature 2014, 510, 273– 277, doi:10.1038/nature13233.

28. Citro, L.; Naidu, S.; Hassan, F.; Kuppusamy, M.L.; Kuppusamy, P.; Angelos, M.G.; Khan, M. Comparison of Human Induced Pluripotent Stem-Cell Derived Cardiomyocytes with Human Mesenchymal Stem Cells following Acute Myocardial Infarction. PLoS One 2014, 9, e116281, doi:10.1371/journal.pone.0116281.

29. Fernandes, S.; Chong, J.J.H.; Paige, S.L.; Iwata, M.; Torok-Storb, B.; Keller, G.; Reinecke, H.; Murry, C.E. Comparison of Human Embryonic Stem Cell-Derived Cardiomyocytes, Cardiovascular Progenitors, and Bone Marrow Mononuclear Cells for Cardiac Repair. Stem Cell Reports 2015, 5, 753–762, doi:10.1016/j.stemcr.2015.09.011.

30. Shiba, Y.; Gomibuchi, T.; Seto, T.; Wada, Y.; Ichimura, H.; Tanaka, Y.; Ogasawara, T.; Okada, K.; Shiba, N.; Sakamoto, K.; et al. Allogeneic transplantation of iPS cell-derived cardiomyocytes regenerates primate hearts. Nature 2016, 538, 388–391, doi:10.1038/nature19815.

31. Weinberger, F.; Breckwoldt, K.; Pecha, S.; Kelly, A.; Geertz, B.; Starbatty, J.; Yorgan, T.; Cheng, K.H.; Lessmann, K.; Stolen, T.; et al. Cardiac repair in Guinea pigs with human engineered heart tissue from induced pluripotent stem cells. Sci. Transl. Med. 2016, 8, 1– 13, doi:10.1126/scitranslmed.aaf8781.

32. Ishida, M.; Miyagawa, S.; Saito, A.; Fukushima, S.; Harada, A.; Ito, E.; Ohashi, F.; Watabe, T.; Hatazawa, J.; Matsuura, K.; et al. Transplantation of Human-induced Pluripotent Stem Cell-derived Cardiomyocytes Is Superior to Somatic Stem Cell Therapy for Restoring Cardiac Function and Oxygen Consumption in a Porcine Model of Myocardial Infarction. Transplantation 2019, 103, 291–298, doi:10.1097/TP.0000000000002384.

33. Romagnuolo, R.; Masoudpour, H.; Porta-Sánchez, A.; Qiang, B.; Barry, J.; Laskary, A.; Qi, X.; Massé, S.; Magtibay, K.; Kawajiri, H.; et al. Human Embryonic Stem Cell-Derived Cardiomyocytes Regenerate the Infarcted Pig Heart but Induce Ventricular Tachyarrhythmias. Stem Cell Reports 2019, 12, 967–981, doi:10.1016/j.stemcr.2019.04.005.

34. Liu, Y.W.; Chen, B.; Yang, X.; Fugate, J.A.; Kalucki, F.A.; Futakuchi-Tsuchida, A.; Couture, L.; Vogel, K.W.; Astley, C.A.; Baldessari, A.; et al. Human embryonic stem cell-derived cardiomyocytes restore function in infarcted hearts of non-human primates. Nat. Biotechnol. 2018, 36, 597–605, doi:10.1038/nbt.4162.

35. Dariolli, R.; Bassaneze, V.; Nakamuta, J.S.; Omae, S.V.; Campos, L.C.G.; Krieger, J.E. Porcine Adipose Tissue-Derived Mesenchymal Stem Cells Retain Their Proliferative Characteristics, Senescence, Karyotype and Plasticity after Long-Term Cryopreservation. PLoS One 2013, 8, e67939, doi:10.1371/journal.pone.0067939.

36. Dariolli, R.; Takimura, C.K.; Campos, C. a; Lemos, P. a; Krieger, J.E. Development of a closed-artery catheter-based myocardial infarction in pigs using sponge and lidocaine hydrochloride infusion to prevent irreversible ventricular fibrillation. Physiol. Rep. 2014, 2, 1–11, doi:10.14814/phy2.12121.

37. Cruvinel, E.; Ogusuku, I.; Cerioni, R.; Rodrigues, S.; Gonçalves, J.; Góes, M.E.; Alvim, J.M.; Silva, A.C.; Lino, V. de S.; Boccardo, E.; et al. Long-term single-cell passaging of human iPSC fully supports pluripotency and high-efficient trilineage differentiation capacity. SAGE Open Med. 2020, 8, 205031212096645, doi:10.1177/2050312120966456.

38. Olivares, E.L.; Ribeiro, V.P.; Werneck de Castro, J.P.S.; Ribeiro, K.C.; Mattos, E.C.; Goldenberg, R.C.S.; Mill, J.G.; Dohmann, H.F.; dos Santos, R.R.; de Carvalho, A.C.C.; et al. Bone marrow stromal cells improve cardiac performance in healed infarcted rat hearts. Am. J. Physiol. Circ. Physiol. 2004, 287, H464–H470, doi:10.1152/ajpheart.01141.2003.

39. Irion, C.I.; Martins, E.L.; Christie, M.L.A.; de Andrade, C.B. V.; de Moraes, A.C.N.; Ferreira, R.P.; Pimentel, C.F.; Suhett, G.D.; de Carvalho, A.C.C.; Lindoso, R.S.; et al. Acute Myocardial Infarction Reduces Respiration in Rat Cardiac Fibers, despite Adipose Tissue Mesenchymal Stromal Cell Transplant. Stem Cells Int. 2020, 2020, 1–19, doi:10.1155/2020/4327965.

40. Fidelis-De-Oliveira, P.; Werneck-De-Castro, J.P.S.; Pinho-Ribeiro, V.; Shalom, B.C.M.; Nascimento-Silva, J.H.; Souza, R.H.C. e; Cruz, I.S.; Rangel, R.R.; Goldenberg, R.C.S.; Campos-De-Carvalho, A.C. Soluble Factors from Multipotent Mesenchymal Stromal Cells have Antinecrotic Effect on Cardiomyocytes in Vitro and Improve Cardiac Function in Infarcted Rat Hearts. Cell Transplant. 2012, 21, 1011–1021, doi:10.3727/096368911X623916.

41. Bartunek, J.; Terzic, A.; Davison, B.A.; Filippatos, G.S.; Radovanovic, S.; Beleslin, B.; Merkely, B.; Musialek, P.; Wojakowski, W.; Andreka, P.; et al. Cardiopoietic cell therapy for advanced ischemic heart failure: results at 39 weeks of the prospective, randomized, double blind, sham-controlled CHART-1 clinical trial. Eur. Heart J. 2016, 38, ehw543, doi:10.1093/eurheartj/ehw543.

42. Chien, K.R.; Frisén, J.; Fritsche-Danielson, R.; Melton, D.A.; Murry, C.E.; Weissman, I.L. Regenerating the field of cardiovascular cell therapy. Nat. Biotechnol. 2019, 37, 232– 237.

43. Cohn, J.N.; Ferrari, R.; Sharpe, N. Cardiac remodeling—concepts and clinical implications: a consensus paper from an international forum on cardiac remodeling. J. Am. Coll. Cardiol. 2000, 35, 569–582, doi:10.1016/S0735-1097(99)00630-0.

44. Giacca, M. Cardiac Regeneration After Myocardial Infarction: an Approachable Goal. Curr. Cardiol. Rep. 2020, 22, 1–8.

45. Leone, M.; Engel, F.B. Advances in heart regeneration based on cardiomyocyte proliferation and regenerative potential of binucleated cardiomyocytes and polyploidization. Clin. Sci. 2019, 133, 1229–1253.

46. Malliaras, K.; Li, T.-S.; Luthringer, D.; Terrovitis, J.; Cheng, K.; Chakravarty, T.; Galang, G.; Zhang, Y.; Schoenhoff, F.; Van Eyk, J.; et al. Safety and efficacy of allogeneic cell therapy in infarcted rats transplanted with mismatched cardiosphere-derived cells. Circulation 2012, 125, 100–12, doi:10.1161/CIRCULATIONAHA.111.042598.

47. Samak, M.; Hinkel, R. Stem Cells in Cardiovascular Medicine: Historical Overview and Future Prospects. Cells 2019, 8, 1530, doi:10.3390/cells8121530.

48. Liang, J.; Huang, W.; Jiang, L.; Paul, C.; Li, X.; Wang, Y. Concise Review: Reduction of Adverse Cardiac Scarring Facilitates Pluripotent Stem Cell-Based Therapy for Myocardial Infarction. Stem Cells 2019, 37, 844–854, doi:10.1002/stem.3009.

49. Jackson, A.O.; Rahman, G.A.; Yin, K.; Long, S. Enhancing Matured Stem-Cardiac Cell Generation and Transplantation: A Novel Strategy for Heart Failure Therapy. J. Cardiovasc. Transl. Res. 2020, 1–17.

50. Fan, C.; Fast, V.G.; Tang, Y.; Zhao, M.; Turner, J.F.; Krishnamurthy, P.; Rogers, J.M.; Valarmathi, M.T.; Yang, J.; Zhu, W.; et al. Cardiomyocytes from CCND2-overexpressing human induced-pluripotent stem cells repopulate the myocardial scar in mice: A 6-month study. J. Mol. Cell. Cardiol. 2019, 137, 25–33, doi:10.1016/j.yjmcc.2019.09.011.

51. Saludas, L.; Garbayo, E.; Mazo, M.; Pelacho, B.; Abizanda, G.; Iglesias-Garcia, O.; Raya, A.; Prósper, F.; Blanco-Prieto, M.J. Long-term engraftment of human cardiomyocytes combined with biodegradable microparticles induces heart repair. J. Pharmacol. Exp. Ther. 2019, 370, 761–771, doi:10.1124/jpet.118.256065.

52. Prabhu, S.D.; Frangogiannis, N.G. The biological basis for cardiac repair after myocardial infarction. Circ. Res. 2016, 119, 91–112.

53. Ferrini, A.; Stevens, M.M.; Sattler, S.; Rosenthal, N. Toward Regeneration of the Heart: Bioengineering Strategies for Immunomodulation. Front. Cardiovasc. Med. 2019, 6, 26.

54. Liehn, E.A.; Postea, O.; Curaj, A.; Marx, N. Repair after myocardial infarction, between fantasy and reality: The role of chemokines. J. Am. Coll. Cardiol. 2011, 58, 2357–2362.

55. Zhao, M.; Fan, C.; Ernst, P.J.; Tang, Y.; Zhu, H.; Mattapally, S.; Oduk, Y.; Borovjagin, A. V; Zhou, L.; Zhang, J.; et al. Y-27632 preconditioning enhances transplantation of human-induced pluripotent stem cell-derived cardiomyocytes in myocardial infarction mice. Cardiovasc. Res. 2019, 115, 343–356, doi:10.1093/cvr/cvy207.

56. Miller, L.W. Cardiovascular Toxicities of Immunosuppressive Agents. Am. J. Transplant. 2002, 2, 807–818, doi:10.1034/j.1600-6143.2002.20902.x.

57. Tavares, P.; Reis, F.; Ribeiro, C.A.F.; Teixeira, F. Cardiovascular effects of cyclosporin treatment in an experimental model. Rev. Port. Cardiol. 2002, 21, 141–55.

58. Zhu, K.; Wu, Q.; Ni, C.; Zhang, P.; Zhong, Z.; Wu, Y.; Wang, Y.; Xu, Y.; Kong, M.; Cheng, H.; et al. Lack of Remuscularization Following Transplantation of Human Embryonic Stem Cell-Derived Cardiovascular Progenitor Cells in Infarcted Nonhuman Primates. Circ. Res. 2018, 122, 958–969, doi:10.1161/CIRCRESAHA.117.311578.

59. Fang, Y.H.; Wang, S.P.H.; Chang, H.Y.; Yang, P.J.; Liu, P.Y.; Liu, Y.W. Immunogenicity in stem cell therapy for cardiac regeneration. Acta Cardiol. Sin. 2020, 36, 588–594.

60. Percie du Sert, N.; Hurst, V.; Ahluwalia, A.; Alam, S.; Avey, M.T.; Baker, M.; Browne, W.J.; Clark, A.; Cuthill, I.C.; Dirnagl, U.; et al. The ARRIVE guidelines 2.0: Updated guidelines for reporting animal research. PLOS Biol. 2020, 18, e3000410, doi:10.1371/journal.pbio.3000410.

61. Koehler, J.; Kuehnel, T.; Kees, F.; Hoecherl, K.; Grobecker, H.F. Comparison of bioavailability and metabolism with two commercial formulations of cyclosporine a in rats. Drug Metab. Dispos. 2002, 30, 658–62.

62. Diehl, R.; Ferrara, F.; Müller, C.; Dreyer, A.Y.; McLeod, D.D.; Fricke, S.; Boltze, J. Immunosuppression for in vivo research: state-of-the-art protocols and experimental approaches. Cell. Mol. Immunol. 2017, 14, 146–179, doi:10.1038/cmi.2016.39.

63. Midha, R.; Mackinnon, S.E.; Evans, P.J.; Best, T.J.; Wong, P.Y. Subcutaneous injection of oral cyclosporin A solution. Microsurgery 1992, 13, 92–4.

64. Sato, Y.; Araki, H.; Kato, J.; Nakamura, K.; Kawano, Y.; Kobune, M.; Sato, T.; Miyanishi, K.; Takayama, T.; Takahashi, M.; et al. Human mesenchymal stem cells xenografted directly to rat liver are differentiated into human hepatocytes without fusion. Blood 2005, 106, 756–63, doi:10.1182/blood-2005-02-0572.

65. Jansen Of Lorkeers, S.J.; Hart, E.; Tang, X.L.; Chamuleau, M.E.D.; Doevendans, P.A.; Bolli, R.; Chamuleau, S.A.J. Cyclosporin in cell therapy for cardiac regeneration. J. Cardiovasc. Transl. Res. 2014, 7, 475–82, doi:10.1007/s12265-014-9570-8.

66. Lang, R.M.; Bierig, M.; Devereux, R.B.; Flachskampf, F. a; Foster, E.; Pellikka, P. a; Picard, M.H.; Roman, M.J.; Seward, J.; Shanewise, J.S.; et al. Recommendations for chamber quantification: a report from the American Society of Echocardiography’s Guidelines and Standards Committee and the Chamber Quantification Writing Group, developed in conjunction with the European Association of Echocardiograph. J. Am. Soc. Echocardiogr. 2005, 18, 1440–63, doi:10.1016/j.echo.2005.10.005.

