## Supplemental File for "*In situ* maturated early-stage human induced pluripotent stem cell-derived cardiomyocytes improve cardiac function by enhancing segmentary contraction in infarcted rats"

**Author's list:**

Diogo Biagi [1], Evelyn Thais Fantozzi [1, #], Julliana C Campos-Oliveira [1, #], Marcus Vinicius Naghetini [1, #], Antonio F. Ribeiro Jr. [1, #], Sirlene Rodrigues [1], Isabella Ogusuku [1, 3], Rubia Vanderlinde [1], Michelle Lopes Araújo Christie [2], Debora B. Mello [2], Antonio C. Campos de Carvalho [2], Marcos Valadares [1], Estela Cruvinel [1] and Rafael Dariolli [1, 4, *]

**Affiliations:**

*[1]PluriCell Biotech, 05508-000, São Paulo, SP, Brazil*

*[2]Carlos Chagas Filho Institute of Biophysics, Federal University of Rio de Janeiro, Rio de Janeiro, 21941-902, Brazil.*

*[3]Gene Center and Department of Biochemistry, Ludwig-Maximilians-Universität München, 81377, München, Germany*

*[4]Department of Pharmacological Sciences, Icahn School of Medicine at Mount Sinai, 10029, New York, NY, USA*

*# These authors equally contributed to this work*

***

### ***Supplementary Experimental Procedures***

**Cardiomyocytes derived from human induced pluripotent stem cells (hiPSC-CM)**

ACP5 hiPSC were expanded in feeder-free conditions using Essencial 8 medium (E8; Thermo Fisher Scientific, USA) until they reached 60–70% confluence. Cells were dissociated and plated (2.5 x 10^5 cells/cm$^2$) with a 3:1 (vol/vol) mixture of E8 and StemFlex media (Thermo Fisher Scientific, USA) supplemented with 10$\mu$M Y-27632 (Cayman Chemical, USA) After 2 days, at day 0 of differentiation, medium was changed to RPMI supplemented with 0.25X B27 supplement (Thermo Fisher Scientific, USA) without insulin (RB-) and 3$\mu$M CHIR99021 (Merck Millipore Sigma, USA). 24 hours later, 50% of the medium was changed to RB- supplemented with 20ng/mL BMP4 (R&D Systems, USA). In day 2 and 4, medium was changed to fresh RPMI supplemented with 0.25X B27 supplement (Thermo Fisher, USA) (RB+) supplemented with 2uM KY2111 and 2uM XAV939 (both from Cayman Chemical, USA). At day 5 and 7, medium was changed to fresh RPMI supplemented with 213 µg/mL Ascorbic Acid (Sigma-Aldrich, USA), 500µg/mL BSA (CDM3). At day 8, cells were cultivated with RPMI without glucose (Thermo Fisher Scientific, USA) for three days. At day 11, media was changed to CDM3 and cells were cultivated until the day before the passage for injection, when they went through the Heat Shock process.

During Heat Shock procedure cultures were subjected to a 30min exposure to 43°C in medium RB+ supplemented with 100 ng/mL IGF-1 (PeproTech; USA) and 0.2 uM cyclosporine A (Sigma-Aldrich; USA), followed by a return to a 37°C medium. Twenty-four hours later, cells were enzymatically isolated with 0.25U/mL Dispase and 0.5mg/mL collagenase II (Both from Sigma-Aldrich, USA) for 1 hour. The membranes were collected centrifuged and further digested with Accutase (Thermo Fisher Scientific, USA). After dilution in 1x HBSS with 1% Fetal Bovine

Serum (Thermo Fisher Scientific, USA) cells were counted and aliquoted in tubes with 10 million cells and kept at 10°C until the time of the syringe preparation.

**Flow cytometry**

Cells were fixed in 4% PFA (LabSynth, Brazil) and kept in the refrigerator for 3-7 days. Protein expression was analyzed by Flow Cytometry (FC) using antibodies for TNNT2, NKX2.5, MLC2V, MLC2A, TNNI1 and TNNI3 listed in Table S1. Data was acquired using CantoII BD equipment and analyzed by FlowJo Software considering 1-2% of false positive events.

**Immunocytochemistry**

iPSC-CM were plated in 96-well plates coated with GELTREX for immunostaing. Cells were also fixed with 4% PFA (LabSynth, Brazil) and permeabilized with Triton 0.1% and Saponin 0.1% (both from Sigma-Aldrich, USA). Next, cells were stained with antibodies for TNNT2, NKX2.5, MLC2V, MLC2A listed in Table S1. All images were generated in EVOS FL (Thermo Fisher Scientific, USA).

**Permanent coronary artery occlusion – myocardial infarction- Induction**

Acute myocardial infarction was induced by permanent occlusion of the left anterior descending coronary artery as described previously with slight modifications [1]. Adult female Wistar rats weighing 150-200 grams were used. After anesthetized these animals were intubated by a cricothyroidotomy procedure and coupled to a mechanical respirator (tidal volume of 2.5mL, respiratory rate of 90-100 irpm). The animals underwent the following surgical procedure: skin incision at the left parasternal level, approximately 1cm long, located 1cm from the mid sternal line, at the junction of the lower and middle thirds between the collarbone and the costal margin. Then, the pectoral muscles, larger and smaller, were dissected aiming at the visualization of the left costal grating. At this time, a purse-string suture of the skin and muscles of the region was made, leaving the knot open until the end of the surgery. With the aid of hemostatic forceps, the incision was made between the 4th and 5th left intercostal space, where the heart was then externalized through the passage and traction of 7-0 prolene thread through the apical region. At this point, the pericardium surrounding the heart was excised. So, the left coronary artery (usually under the left atrium) was permanently occluded with 7-0 prolene thread as close as possible to its origin in the aorta. Then the heart was quickly repositioned to its original anatomical site and the pouch suture knot finally tightened. The groups were randomized after induction of myocardial infarction. The sham operated group (SHAM) underwent the same surgical stress, but without coronary artery occlusion.

**Echocardiographic assessments**

In order to assess cardiac function, all the animals in this study were subjected to echocardiogralhy pre- and post-treatments (On days 6 and 37 after MI-induction). Parasternal-long and short axis images were captured using VEVO 2100 ultrasound equipment (VisualSonics Vevo® 2100 Imaging System, Canada) with a 25 MHz transducer. Analyzes were performed *off-line* using VisualSonics software (VevoCQ™ LV Analysis - VisualSonics, Canada). Parameters such as Systolic and Diastolic volumes, were calculated using *Modified Simpson Algorithm* present in the Analysis software (parasternal-short axis sequential imaging from base to apex). Based on these volumes, the left ventricle ejection fraction (LVEF, %) was calculated pre- and post-injection treatments (On days 6 and 37 after MI induction). Also, linear measurements were obtained from parasternal-short and -long axis images. Left ventricle shortening fraction (LVSF, %) was calculated, using systolic and diastolic diameters. Still, left ventricle mass (LV mass, mg) was estimated by linear measurements. Beating rate (beats per minute – BPM) was calculated directly by animal table-ECG system connected to VEVO 2100 system. Echocardiographic results were interpreted considering the American Society of Echocardiography recommendations [2] concerning the rat model. All parameters were shown as the mean values of three consecutive cardiac cycles. Transthoracic echocardiography image acquisition and analysis was performed by 2 experts blinded to the experimental groups.

In addition, Speckle-Tracking analyses to access LV radial, circumferential and longitudinal strains were performed using B-mode short- and long-axis 300 frames images using Vevo Strain Analysis Software (Vevo LAB - VisualSonics, Canada). The analysis was performed following the manufacture's user guide. Mean Strain Time-to-Peak (AVG PK %) was used as one of the proxies of strain. This parameter was assessed on both short- and long- axis images for radial and circumferential, and radial and longitudinal strain respectively. Also, Global Circumferential Strain (GCS) and Global Longitudinal Strain (GLS) were calculated from short- and long-axis images, respectively. Short-axis analyses were performed at papillary-level and long-axis analyses at the medial portion of the parasternal long-axis image (where the entire LV can be observed in the presence of the mitral and aortic valves). All Strain analyses incorporated 3-5 consecutive cardiac cycles. Algorithm and equation details can be found in the manufacture's user guide.

**LVEF pre-injection-based exclusion Criteria**

Six days after MI induction rats were subjected to an echocardiographic assessment of cardiac. An on-line analysis of the LVEF was performed by a technician and these results were used to randomize MI induced rats in two groups, namely: PSC (placebo for cells that received injections of vehicle – Pro-survival cocktail) and CELL (PSC (vehicle) + 10 million hiPSC-CMs). SHAM and CTRL animals were also imaged.

To investigate heart function improvement after cell injection into infarcted rats, we excluded from the analysis animals that had shown less than 20% decrease on LVEF when compared to control animals (or up to 55% LVEF) indicating functional compromise after infarction. Due to the statistical difference observed in the randomization procedure, indicating high level of heterogeneity in the injected group, we excluded animals of the cell injected group with LVEF below the minimum value observed in the PSC group, effectively leveling the two groups for direct comparison.

**Anatomopathological injury assessments**

Some measurements were determined by picrosirius red stain. This procedure consists in three steps, first the slides were deparaffinized with xylol (Êxodo Cientifica, Brazil) followed by a decreasing concentration battery of ethanol (Cirurgica Estilo, Brazil) solution washes. Then, slides were stained for 20min in Bouin fixative solution (Êxodo Científica, Brazil), washed in distillated water and finally a last bath with 0.1% Sirius red (Êxodo Científica, Brazil) in aqueous saturated picric acid solution (Imbralab, Brazil) was performed. Next, the slides were finalized using aqueous mount media (Vector, USA). To quantify injury perimeter and wall thickness on left ventricle ImageJ software was used as shown in ***Figure***. By the software we have measured the injury and healthy perimeters using freehand selections tool to calculate the percentage of injury (***Figure A***). For the wall thickness values we have calculated the mean value of three different measurements with straight line tool from the injury and healthy areas (***Figure B***). In the same picrosirius stained slides, we also evaluated MI area and interstitial collagen using Image-Pro-Plus Software. To evaluate the area of injury, we first remove the right ventricle from the scanned 4x images with image editor and then, we measured the percentage of injury by selecting the colors red (injury) and yellow (healthy) using the Count/Size tool (***Figure C***). Similar procedure was performed to measure interstitial collagen from 20 images taken at 20x in the injury and remote areas.

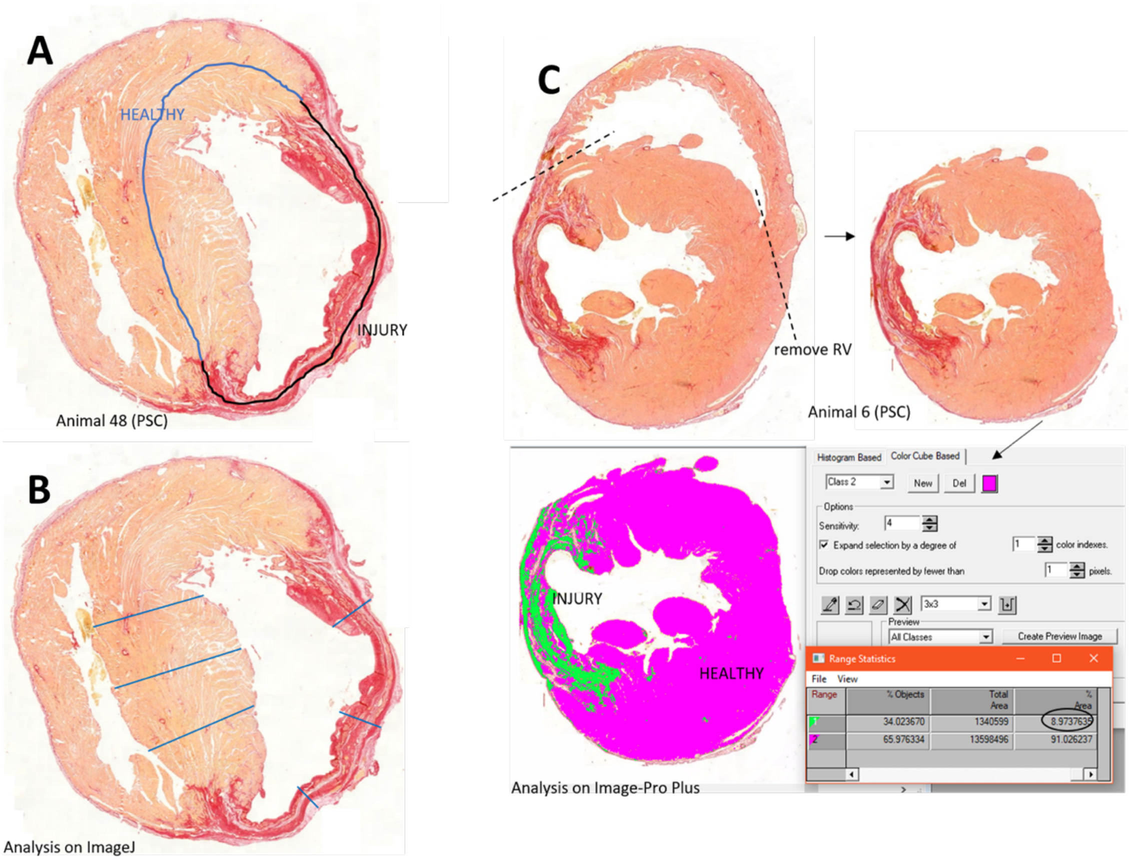

**Figure:** *Examples of picrosirius red stain analysis*. In A: perimeter area. In B: wall thickness. In C: injury area.

**Human Ku80 positive graft assessments**

To assess human graft tissues into the LV of injected rats, the sections were deparaffinized, then, heat-induced epitope retrieval with electric pressure cooker for 15min, quenched for unspecific binding with 5% donkey serum (Sigma, USA) + 0.5% TWEEN 20 in PBS (PBST) for 45min and incubated overnight at 4°C with primary antibody (antibodies list – Table S1). After 16 hours slides were washed with PBST and incubated with a secondary antibody (1:500) for 1h.

In IF, the reaction was used autofluorescence quenching kit (Trueview - Vector, USA). Finally, the slides were finalized using aqueous mount VectaShield media (Vector, USA) with DAPI. In IHC, the reaction was amplified with avidin-biotin complex (Vector, USA) and reveled with DAB (Vector, USA). Counterstain was made with hematoxylin (Êxodo Cientifica, Brazil) for 5min. After immunostaining, the slides were finalized using aqueous mount media (Vector, USA). Immunofluorescence and immunohistochemistry micrographs were obtained using TissueFAXS slide scanner (TissueGnostics, Austria), confocal microscope LSM 800 (Zeiss-Germany) and a conventional fluorescence microscope Axio Imager 2 (Zeiss-Germany). The analyses were performed using ImageJ through the multi-point tool used to count positive markers and nuclei and calculate the percentage of positive events.

**Biodistribution**

DNA were isolated from liver, brain, lung, kidney, and spleen cryopreserved in -80°C from all animals using Wizard™ Genomic DNA Purification Kit (Promega, USA) following manufacturer's instructions. Then, end-point PCR were run with 200ng of all samples targeting human mitochondria: forward primer 5′-CACCGGCGCAGTCATTCTCATA-3′ and reverse primer 5′-GAGTCCTGTAAGTAGGAGA-3′ (265bp) [3]. Positive control reactions were done using 200ng of rat DNA supplemented with 10ng, 1ng, 100pg, 10pg, and 1pg of human DNA. Water was used as negative control and a rat specific Ahr sequence were also amplified: forward primer 5′-TGCCAGCAACAGCCTGTGAG-3′ and reverse primer 5′-AACTGGCGAACATGCCATTGA-3′ (650bp) [4]. PCR were analyzed by agarose gel electrophoresis (2% w/v).

All tissues that were positive for human DNA had 2 more DNA isolation from different sites from the same tissue sample. Moreover, quantitative PCR were run targeting mitochondrial sequence as described above. PCR were run in a QuantiStudio 3 Real-time system from Applied Biosystems. A standard curve of 200ng of rat DNA supplemented with 10ng, 1ng, 100pg, 10pg, and 1pg of human DNA were used and all sample were run in triplicates. Considering that a human cell DNA has 5pg [5], this assay can detect 1 human cell in 200.000 cells.

**References**

1. Olivares, E.L.; Ribeiro, V.P.; Werneck de Castro, J.P.S.; Ribeiro, K.C.; Mattos, E.C.; Goldenberg, R.C.S.; Mill, J.G.; Dohmann, H.F.; dos Santos, R.R.; de Carvalho, A.C.C.; et al. Bone marrow stromal cells improve cardiac performance in healed infarcted rat hearts. *Am. J. Physiol. Circ. Physiol.* **2004**, *287*, H464–H470, doi:10.1152/ajpheart.01141.2003.
2. Lang, R.M.; Bierig, M.; Devereux, R.B.; Flachskampf, F. a; Foster, E.; Pellikka, P. a; Picard, M.H.; Roman, M.J.; Seward, J.; Shanewise, J.S.; et al. Recommendations for chamber quantification: a report from the American Society of Echocardiography's Guidelines and Standards Committee and the Chamber Quantification Writing Group, developed in conjunction with the European Association of Echocardiograph. *J. Am. Soc. Echocardiogr.* **2005**, *18*, 1440–63, doi:10.1016/j.echo.2005.10.005.
3. Liu, Y.W.; Chen, B.; Yang, X.; Fugate, J.A.; Kalucki, F.A.; Futakuchi-Tsuchida, A.; Couture, L.; Vogel, K.W.; Astley, C.A.; Baldessari, A.; et al. Human embryonic stem cell-derived cardiomyocytes restore function in infarcted hearts of non-human primates. *Nat. Biotechnol.* **2018**, *36*, 597–605, doi:10.1038/nbt.4162.
4. Adamovic, T.; McAllister, D.; Guryev, V.; Wang, X.; Andrae, J.W.; Cuppen, E.; Jacob, H.J.; Sugg, S.L. Microalterations of Inherently Unstable Genomic Regions in Rat Mammary Carcinomas as Revealed by Long Oligonucleotide Array-Based Comparative Genomic Hybridization. *Cancer Res.* **2009**, *69*, 5159–5167, doi:10.1158/0008-5472.CAN-08-4038.
5. Sambrook, J. *Molecular cloning : a laboratory manual*; Third edition. Cold Spring Harbor, N.Y. : Cold Spring Harbor Laboratory Press, [2001] ©2001;

### ***Supplementary Tables***

**Table S1:** LVEF distribution in MI-induced rats after randomization.

| Group | Min (%) | Max (%) | Mean (%) | STD |
|---|---|---|---|---|
| PSC | 38.0 | 51.6 | 44.0 | 5.5 |
| CELL | 38.3 | 52.6 | 43.7 | 5.8 |

**Table S2:** Percentage of hKu80 positive versus scar tissue area per animal.

| Animal ID | $hKu80^+$ ($mm^2$) | Injury ($mm^2$) | % of human graft |
|---|---|---|---|
| **15** | 0.18 | 4.78 | 3.62% |
| **21** | 0.65 | 6.30 | 9.28% |
| **29** | 0.99 | 3.96 | 19.93% |
| **31** | 0.09 | 2.14 | 3.90% |
| **39** | 0.14 | 9.50 | 1.44% |
| **45** | 0.15 | 6.54 | 2.30% |
| **49** | 0.14 | 4.21 | 3.12% |

**Table S3:** Antibodies list for flow cytometry (FC), immunocytochemistry (ICC), immunohistochemistry (IHC) and histological immunofluorescence staining (IF). n/a means no apply.

| Antibody | Procedure used and dilution | Company/catalogue number (#) | Retrieval buffer for IF |
|---|---|---|---|
| TNNI3 | 1:250 (IF) | Abcam/ab52862 | Tris/EDTA pH 9.0 |
| TNNI3 | 1:30 (FC) | DSHB/TI-4 | n/a |
| TNNI1 | 1:500 (IF) | Thermo/701585 | n/a |
| TNNT2 | 1:250 (IF)<br>1:2000 (ICC) | Fitzgerald/70C<br>CR4037GAP | Citrate pH 6.0<br>n/a |
| TNNT2 | 1:500 (FC) | Thermo/MA5-12960 | n/a |
| Ku-80 | 1:100 (IF) | CellSignaling/2180S | Citrate pH 6.0 |
| Ki-67 | 1:100 (IF) | Abcam/ab15580 | Citrate pH 6.0 |
| Phh3 | 1:100 (IF) | Abcam/ab5176 | Citrate pH 6.0 |
| vWF | 1:100 (IF) | Abcam/ab6994 | Citrate pH 6.0 |
| Pan-cadherin | 1:100 (IF) | Sigma/C3678 | Citrate pH 6.0 |
| Caveolin-3 | 1:250 (IF) | Abcam/ab2912 | Citrate pH 6.0 |
| Conexin 43 | 1:1000 (IF) | Abcam/ab11370 | Citrate pH 6.0 |
| CD 03 | 1:150 (IHC) | Abcam/ab16669 | Citrate pH 6.0 |
| CD 20 | 1:250 (IHC) | BioCare/ACR3004 | Citrate pH 6.0 |
| CD45 | 1:250 (IHC) | Abcam/ab10558 | Citrate pH 6.0 |
| CD 31 | 1:400 (IF) | CellSignaling/3528 | Citrate pH 6.0 |
| Cytokeratin | 1:50 (IF) | Dako/M3515 | Citrate pH 6.0 |
| MLC2 a | 1:100 (IF)<br>1:1000 (ICC) | Thermo/MA5-26390 | Tris/EDTA pH 9.0<br>n/a |
| MLC2 a | 1:750 (FC) | SynapticSystems/311 011 | n/a |
| MLC2 v | 1:200 (IF) | BD/565497 | Tris/EDTA pH 9.0 |
| MLC2 v | 1:300 (FC)<br><br>1:500 (ICC) | Abcam/ab79935 | n/a |
| NKX-2.5 | 1:100 (FC and ICC) | CellSignaling/8792S | n/a |

**Table S4:** List of reagents used in this study.

| Reagents | Company | Country | #catalogue |
|---|---|---|---|
| Accutase | Thermo Fischer Scientific | USA | A1110501 |
| Acid Acetic | Êxodo Cientifica | USA | AA09870RA |
| agarose | Thermo Fischer Scientific | USA | 16500500 |
| Alkaline phosphatase polymer kit | Vector | USA | MP-5402 |
| Aniline blue | Êxodo Cientifica | USA | AA09890RA |
| aqueous slide mount solution | Vector | USA | H-5501 |
| Ascorbic Acid | Sigma-Aldrich | USA | 4403 |
| Avidin-biotin complex | Vector | USA | S -2001 |
| B27 | Thermo Fisher Scientific | USA | 17504044 |
| B27 minus insulin | Thermo Fisher Scientific | USA | A1895601 |
| BMP4 | R&D Systems | USA | 341-BP-050 |
| Bouin fixative solution | Êxodo Cientifica | BRA | FB08835SO |
| BSA | Sigma-Aldrich | USA | A2153-100 |
| CHIR99021 | Merck Millipore Sigma | USA | 361571 |
| Collagenase II | Sigma-Aldrich | USA | 17101015 |
| Cyclosporine A | Sigma-Aldrich | USA | 30024 |
| Cyclosporine A Sandimmun | Novartis | SWZ | 945570 |
| DAB | Vector | USA | SK4105 |
| DAPI | Thermo Fischer Scientific | USA | D1306 |
| Dispase | Sigma-Aldrich | USA | D4693 |
| Donkey serum | Millipore | USA | S30 |
| Essencial8 | Thermo Fisher Scientific | USA | A1517001 |
| Ethanol | Cirurgica Estilo | BRA | 13462 |
| Fetal Bovine Serum | Thermo Fisher Scientific | USA | 12657029 |
| GelTREX | Thermo Fischer Scientific | USA | A1413302 |
| Glucose | Sigma-Aldrich | USA | G8270 |
| Hematoxylin | Êxodo Cientifica | USA | HH09490SO |
| Hoechst | Sigma-Aldrich | USA | 14533 |
| IGF-1 | PeproTech | USA | AF-100-11 |
| KY2111 | Cayman Chemical | USA | 14315 |
| PBS | Thermo Fischer Scientific | USA | 21600-044 |
| PFA | LabSynth | BRA | P1005.06.AG |
| Picric acid | Imbralab | BRA | 4283 |
| Red chromagen | Vector | USA | SK5100 |
| RPMI 1640 | Thermo Fisher Scientific | USA | 11875093 |
| Saponin | Sigma-Aldrich | USA | 84310 |
| Sirius red | Êxodo Cientifica | BRA | SR07371RA |
| StemFlex | Thermo Fisher Scientific | USA | A3349301 |
| Triton X-100 | Sigma-Aldrich | USA | T8787 |
| Trueview (Autofluorescence quenching) | Vector | USA | SP-8400 |
| Tween 20 | Sigma-Aldrich | USA | P1379-100 |
| VectaShield | Vector | USA | H1700 |
| Wizard Genomic DNA purification Kit | Promega | USA | A1120 |
| XAV939 | Cayman Chemical | USA | 13596 |
| Xylol | Êxodo Cientifica | BRA | X09709RA |
| Y-27632 | Cayman Chemical | USA | 10005583 |

### ***Supplementary Figures***

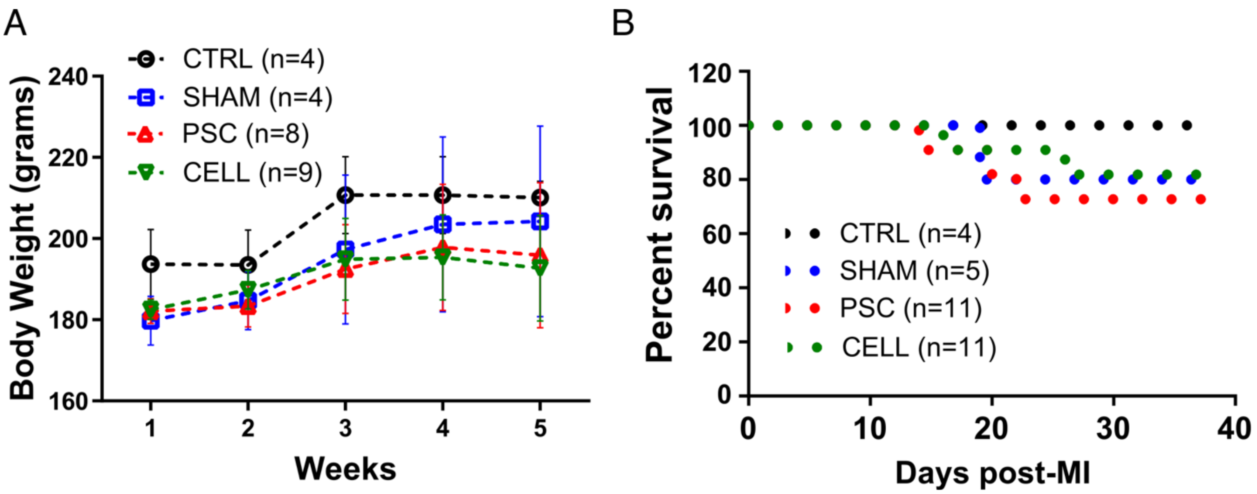

**Figure S1:** *Body weight control and survival rate of CsA immunosuppressed and infarcted rats that completed the experimental protocol*.

**A)** The immunosuppression with Ciclosporin A was weekly adjusted based on the animal's body weight, as well as potential toxicity of the treatment.

**B)** Kaplan-Meier survival curve. We did not observe any significant different in mortality/survival.

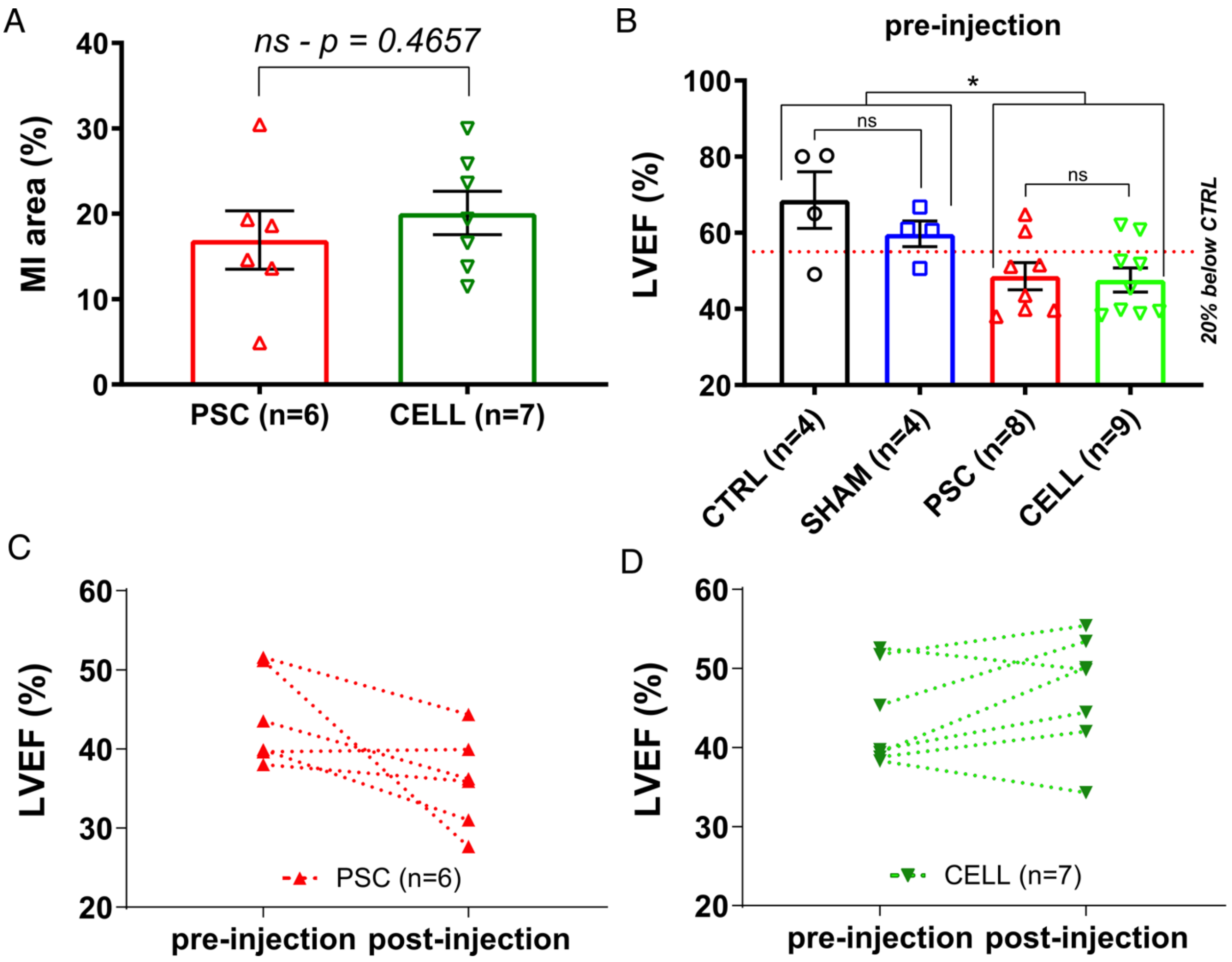

**Figure S2:** *MI area and LVEF pre-injections*. Functional improvements can be associated to the identification of human grafts into the fibrotic scar. Picrosirius red stained representative cross-sectional image were used to quantify the percentage of MI area 30 days after treatment.

**A)** Quantification of MI area (mean ± SE). **B)** Animals that completed the experimental therapy had their LVEF pre-treatment used to exclude rats without significant impairment of cardiac function. Red dashed line represents the pre-treatment limit for animals to be considered in the next steps of analysis. This threshold was calculated based on the mean LVEF of the CTRLT group subtracting 20% of this value to establish the cut-off. Note that 2 rats from PSC group and 2 from CELL group are above the limit. These 4 animals were excluded from the next set of functional and molecular assessments. Pre-injections both couples CTRL and SHAM and PSC and CELL animals displayed similar LVEF. Also, as expected PSC and CELL rats showed deteriorated LVEF compared with CTRL and SHAM animals. $P<0.05$. **C)** LVEF values Pre- and post-injection for each animal from PSC group. **D)** LVEF values Pre- and post-injection for each animal from CELL group.

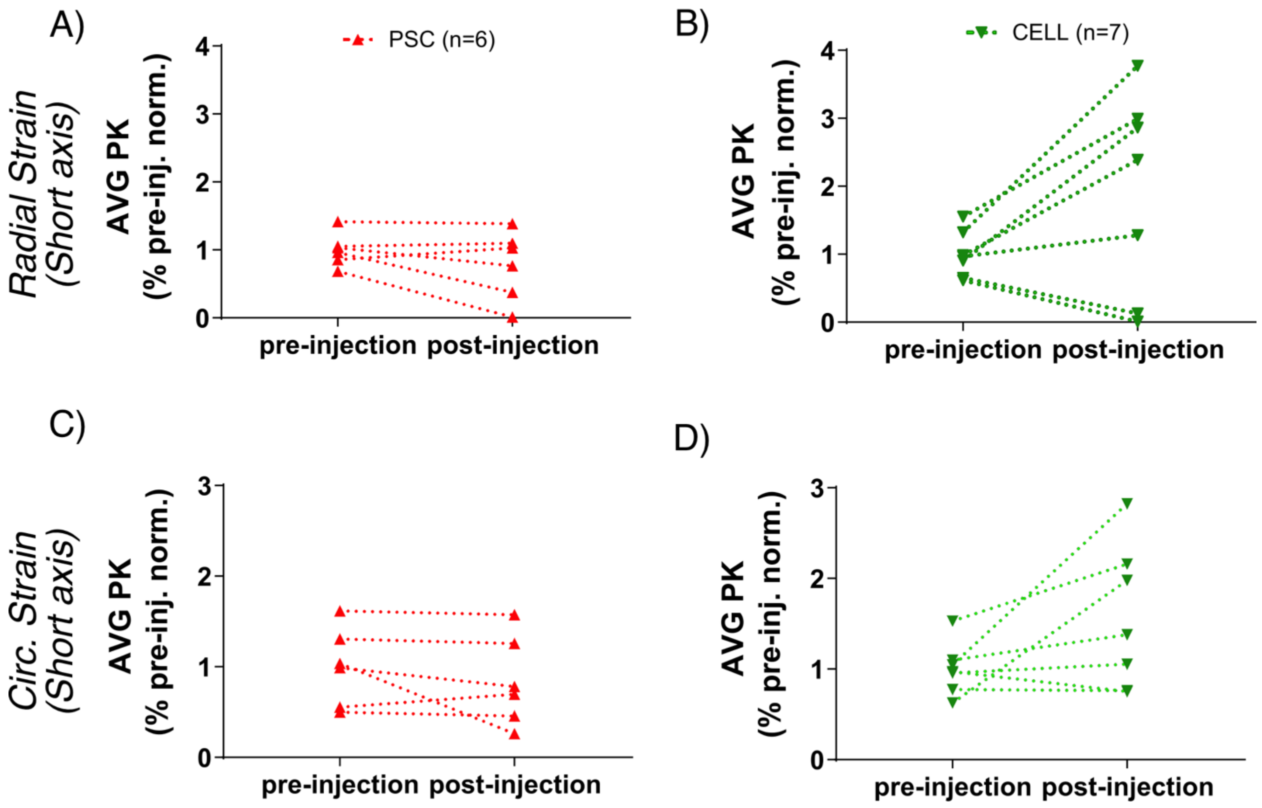

**Figure S3:** *Short Axis Strain Analysis*. Radial and circumferential strain time-to-peak were calculated using papillary level images and the average time-to-peak (AVG PK) for all the six segments normalized by mean pre-injection values for each group to access how treatment modulated this parameter. **A)** Radial Strain time-to-peak (AVG PK) of PSC rats normalized by pre-treatment values. **B)** Radial Strain time-to-peak (AVG PK) of CELL rats normalized by pre-treatment values. **C)** Circumferential Strain time-to-peak (AVG PK) of PSC rats normalized by pre-treatment values. **D)** Circumferential Strain time-to-peak (AVG PK) of CELL rats normalized by pre-treatment values.

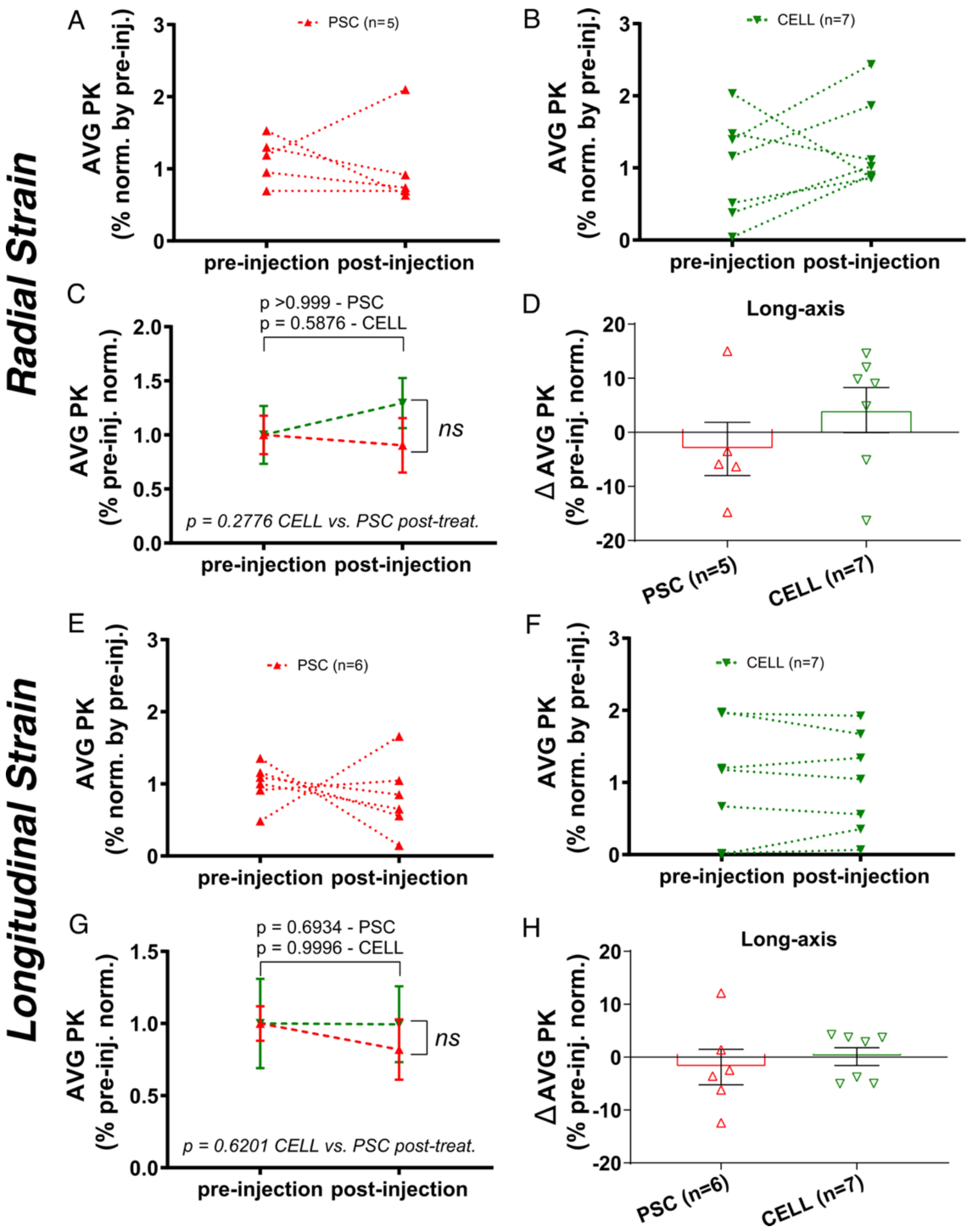

**Figure S4:** *Long Axis Strain Analysis*. Radial and longitudinal strain time-to-peak were calculated using longitudinal mid-portion images and the average time-to-peak (AVG PK) for all the six segments normalized by mean pre-injection values for each group to access how treatment modulated this parameter. **A)** Radial Strain time-to-peak (AVG PK) of PSC rats normalized by pre-treatment values. **B)** Radial Strain time-to-peak (AVG PK) of CELL rats normalized by pre-treatment values. **C)** Mean Radial Strain time-to-peak (AVG PK) of PSC and CELL rats were normalized by their respective pre-treatment values. **D)** Radial Strain time-to-peak delta post- minus pre-injection. **E)** Longitudinal Strain time-to-peak (AVG PK) of PSC rats normalized by pre-treatment values. **F)** Longitudinal Strain time-to-peak (AVG PK) of CELL rats normalized by pre-treatment values. **G)** Mean Longitudinal Strain time-to-peak (AVG PK) of PSC and CELL rats were normalized by their respective pre-treatment values. **H)** Longitudinal Strain time-to-peak delta post- minus pre-injection.

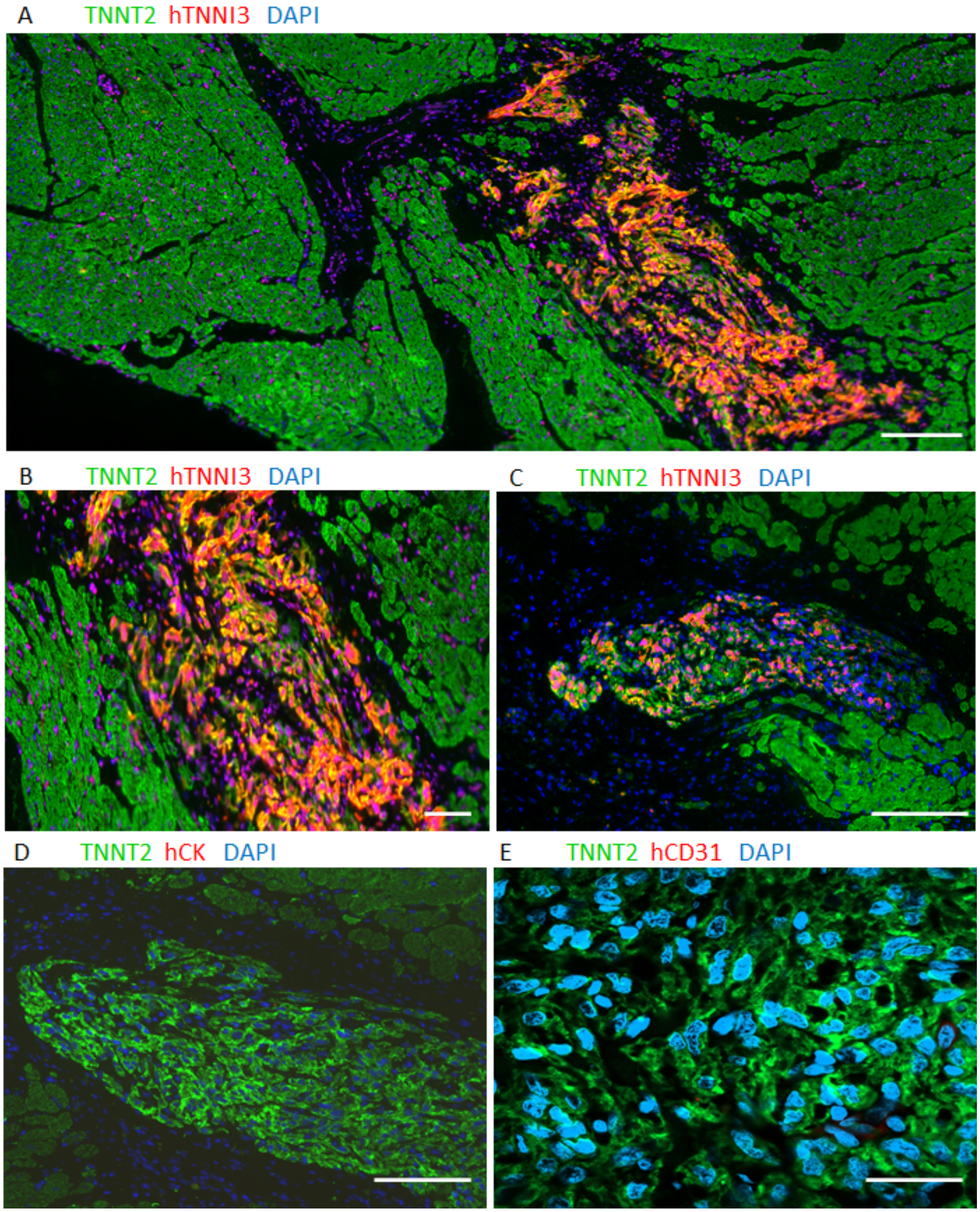

**Figure S5:** *Morphology of the human graft*. **A)** location of the graft in rat tissue marked for hTNNI3 (red), TNNT2 (green) and nuclei (blue). **B)** Magnification of A, showing sarcomeres' organization **C)** Another example of human graft expressed TNNI3. **D)** no red staining for human CK. **E)** non-expression of human endothelial cells positive for CD31. Scale bar: 50 $\mu$m.

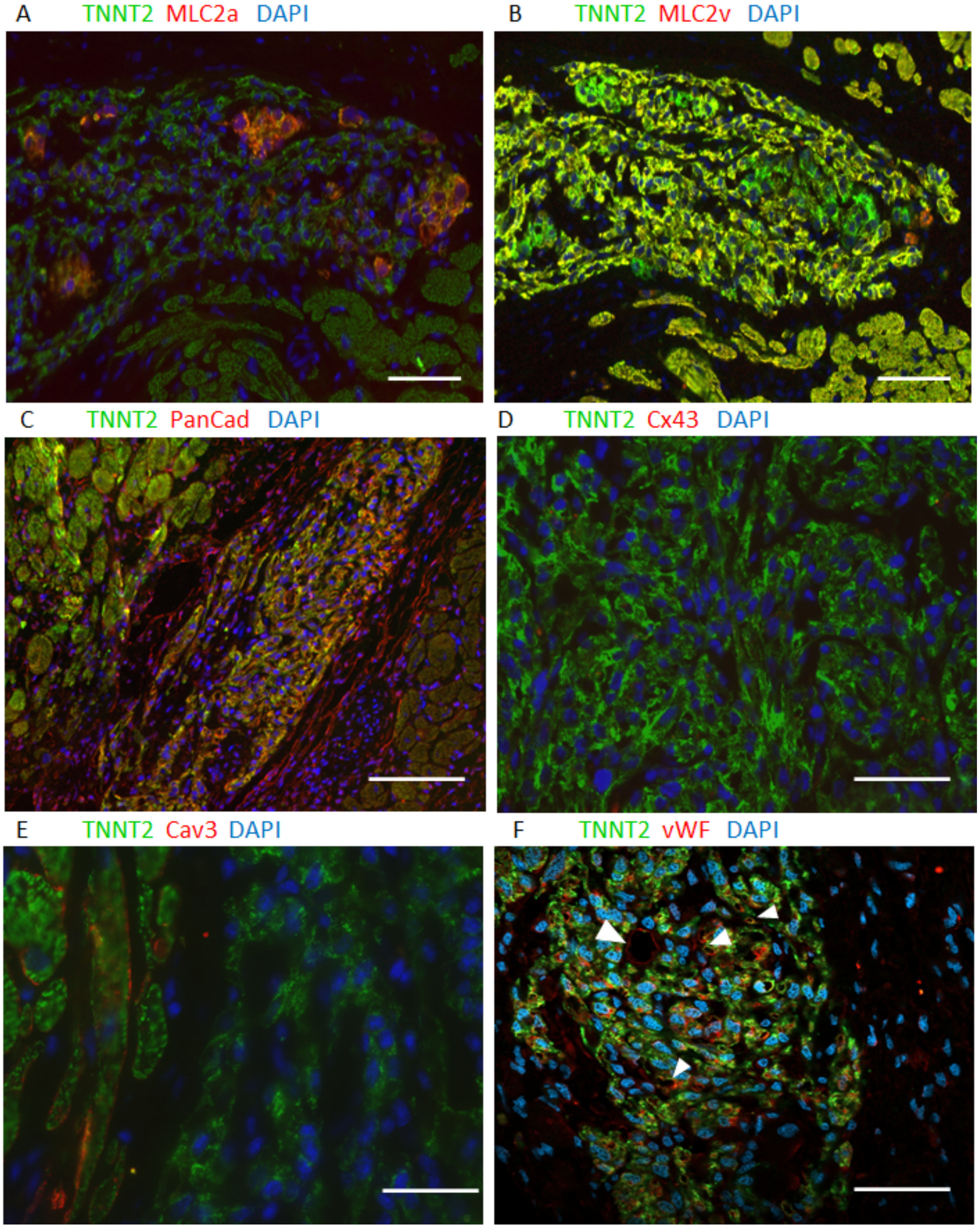

**Figure S6:** *Maturation and cell-cell interaction in the human graft by immunofluorescence staining*. **A)** Costaining for TNNT2 (green) and MLC2a (red), nuclei were counterstaining with DAPI (blue). **B)** Immunostaining for MLC2v and TNNT2 with DAPI in the same graft marked with MLC2a **(A)**. General cardiac marker TNNT2 (green) and red subtype marker PanCad **(C)**, Connexin 43 **(D)**, Caveolin 3 **(E)** and vWF **(F)** with arrowheads show vascularization within in the graft. Scale bar: 50 $\mu$m.

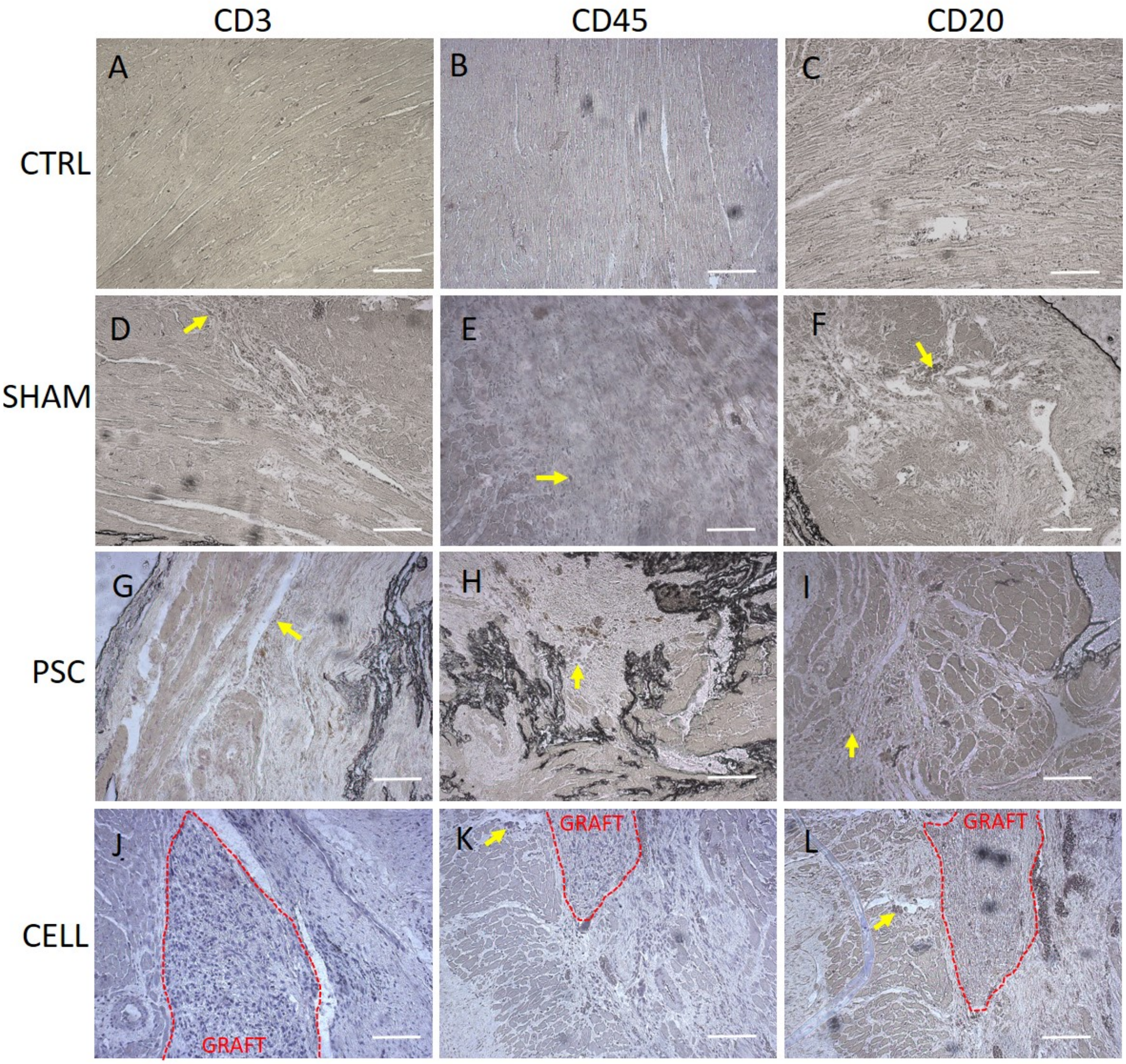

**Figure S7:** *Immunohistochemistry for inflammatory markers*. **A-C** No evidence for CD3, CD45 and CD20 respectively, in healthy control group. **D-F)** Difuse labeling for CD3, CD45 and CD20 in SHAM group. PSC group shows difuse **(G;I)** and patches **(H)** labeling for all markers. Cell group shows no evidence for CD3 labeling **(J)** but shows patches of CD20 and CD45 markers **(K-L)**. Scale bar: 50 $\mu$m.

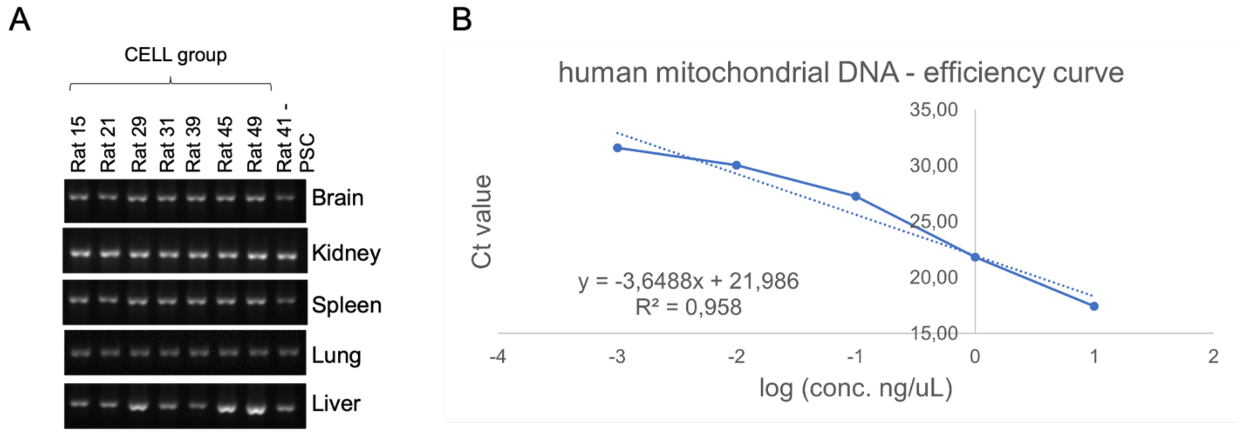

**Figure S8:** *Human mitochondrial DNA were identified in some tissues other than heart and subjected to qPCR analysis*. **A)** Rat DNA sequence was amplified as control for reactions. **B)** Quantitative human mitochondrial DNA. Standard curve of human mitochondrial DNA sequence in 1pg, 10pg, 100pg, 1ng and 10ng of human DNA diluted in 200ng of rat DNA per sample. We considered 1pg of human DNA our lowest detectable amplification.

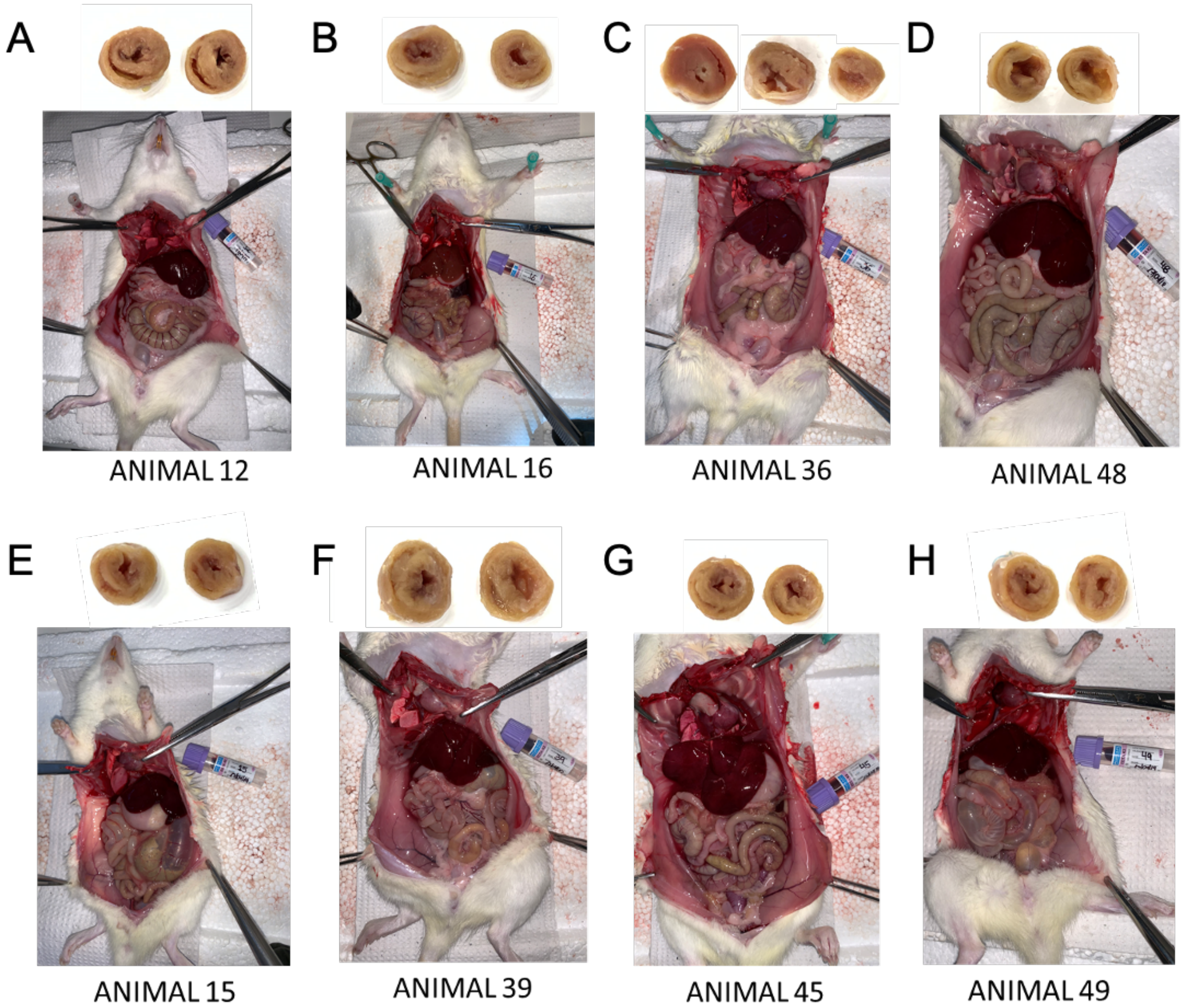

**Figure S9:** *Necropsy*. On day 37, rats underwent deep anesthesia were euthanized by the direct injection of KCl overdose. **A)** Healthy control rat. **B)** SHAM rat. **C and D)** PSC rats (MI-induced animals). **E-F)** CELL-treated rats (MI-induced) whom human DNA was found out of the heart.
